# Shared Latent Decision Strategies Underlie Reward-Guided Behavior Across Species

**DOI:** 10.64898/2026.09.02.748514

**Authors:** Zahra Rostami, Hadi Choubdar Parvin, Eshaan S. Iyer, Vitaro Peter, Rebecca Boehme, Markus Heilig, R Becket Ebitz, Leah M Mayo, Rosemary C Bagot

## Abstract

Adaptive behavior requires that organisms learn which actions are rewarded and to update action selection when the environment changes. While human and non-human animals exhibit adaptive behavior, whether apparently similar behavior reflects common decision strategies remains unclear. Probabilistic reversal learning provides a cross-species assay of reward-guided choice, yet standard metrics such as accuracy or reward rate can obscure underlying strategies that generate choices. Here, we applied parallel probabilistic reversal learning tasks in mice and humans and used a generalized linear model–hidden Markov model to infer latent decision strategies from trial-by-trial behavior. Across species, choices were organized into stable behavioral states with differing reliance on choice history, reward history, and response bias. Among these latent states, we identify a conserved reward-learning strategy in mice and humans characterized by the greatest feedback sensitivity, reward efficiency, and adaptation after reversal. Simulating choice behavior using state-specific decision policies reproduced the empirical hierarchy of performance, confirming that the latent states capture meaningful behavioral strategies. Although mice and humans differ in the temporal dynamics of reward learning, both species ultimately converge on the same optimized strategy. These findings identify a conserved latent reward-learning strategy in mice and humans, defining a translational framework for studying how adaptive decision-making is shaped by task experience, stress, affective processes, and neural circuit function.

**Teaser:** A shared latent behavioral strategy underlies flexible decision-making in mice and humans.

## INTRODUCTION

Reward-guided decision-making is fundamental for adaptive behavior, allowing individuals to use past outcomes to select actions and adjust their behavior as environmental conditions change(*1*). Probabilistic reversal learning (PRL) tasks are widely used to study reward-guided decision-making and cognitive flexibility across species (*2*). In these paradigms, subjects repeatedly choose between two options associated with distinct reward probabilities that change across time (*3*). By requiring continuous adaptation to changing contingencies, PRL tasks provide a controlled framework for examining how behavior shifts between reward-guided and alternative response strategies as environmental demands change (*3*).

PRL tasks have been extensively used to investigate the neural and computational mechanisms underlying adaptive behavior in humans and non-human animal models(*3*). Altered reversal learning has been implicated in multiple neuropsychiatric disorders, including depression, obsessive-compulsive disorder, schizophrenia, Parkinson’s disease, addiction, and neurodevelopmental disorders(*4–6*). In particular, abnormal switching after probabilistic non-reward and impaired adaptation to changing contingencies may reflect dysfunction within frontocorticostriatal circuits underlying reward-guided behavior. Consequently, PRL has emerged as a powerful translational assay for studying impulsivity, compulsivity, reward sensitivity, and cognitive flexibility across species(*4, 5*).

Cross-species approaches are uniquely valuable in identifying evolutionarily conserved behavioral motifs that can be mechanistically interrogated in animal models(*7*). Rodent studies provide access to powerful experimental tools for measuring and manipulating neural activity yet the relevance of these findings to human cognition can be difficult to establish (*8*). Parallel investigations in humans and rodents offer an important strategy for determining the extent to which observable behaviors arise from shared or distinct computational mechanisms(*9*). PRL has been adapted successfully across species, yet task similarity is insufficient to establish conserved computational processes; different species may leverage qualitatively different strategies to solve the same task limiting translational relevance. Indeed, even when PRL tasks are designed to be closely comparable, humans and rodents can differ in persistence, exploration, and sensitivity to probabilistic feedback (*9–11*). Such behavioral differences make it unclear whether PRL engages similar underlying strategies across species or whether comparable task demands are solved through species-specific behavioral policies.

A growing body of work suggests that behavior during reversal learning cannot be fully explained by a single stationary strategy, rather subjects dynamically alternate between multiple latent behavioral modes over time(*3, 9*). However, the extent to which latent strategies are conserved across species remains unclear. Determining whether analogous PRL tasks engage common computational states in humans and rodents is critical to support the translational interpretation of preclinical models.

Here, we implemented parallel PRL paradigms in humans and mice and modeled behavior using a generalized linear model–hidden Markov model (GLM-HMM) framework to reveal discrete latent behavioral states from trial-by-trial choices and capture probabilistic transitions between strategies over time. In humans and mice, we identified conserved reward-sensitive states corresponding to *reward-learning* and *reward-achievement*, as well as additional species-specific strategies. Mice transiently expressed a *choice-persistence* state characterized by perseveration independent of outcome, whereas humans expressed a *reward-history* state integrating recent outcome information. Despite these differences, in both species optimal behavioral performance converged on a shared reward-learning state. Moreover, the emergence of this state followed analogous logistic growth dynamics across trials, although humans transitioned into *reward-learning* substantially faster than mice. Together, these findings reveal a conserved latent structure underlying probabilistic reversal learning across species while highlighting species-specific adaptations in strategy organization and learning dynamics. By identifying a conserved reward-learning state across species, this work paves the way for precise mechanistic translation from rodent models to human cognition.

## RESULTS

### Probabilistic reversal learning engages stable latent strategies in mice

To examine reward-based decision making strategies in mice, we trained mice in a two-lever probabilistic reversal learning (PRL) task in which each lever delivered chocolate milk reward with either high (80%) or low (20%) probability (Fig. 1A) with five consecutive responses on the high-probability lever triggering contingency reversal to maintain environmental volatility. Across 13 training sessions, mice (n = 23; 11 female) progressively completed more reversals while maintaining a relatively stable reward rate (fig. S1), indicating gradual acquisition of task structure across days that was similar across sexes (fig. S1).

**Fig. 1.**
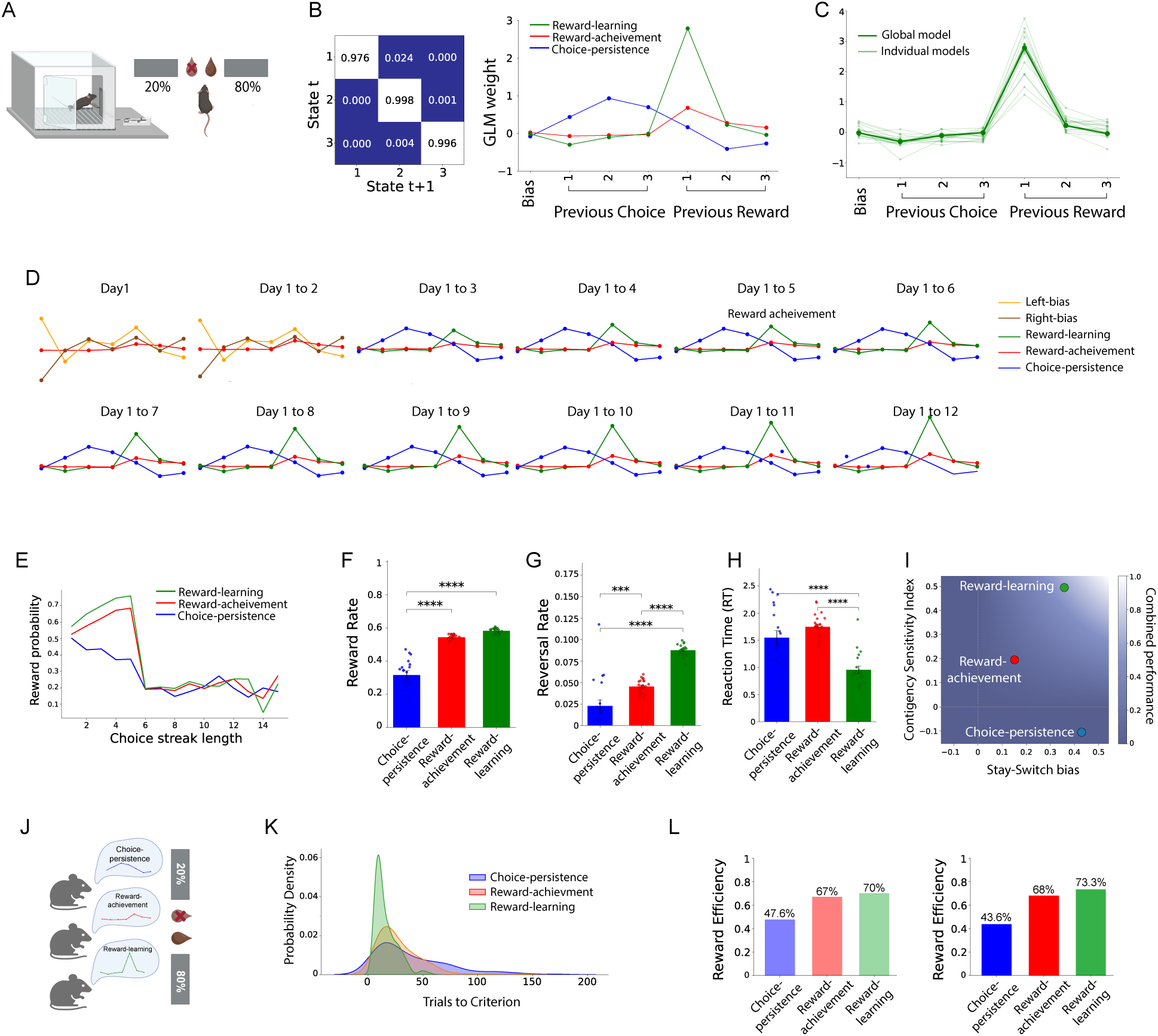
Latent decision strategies govern probabilistic reversal learning in mice. **(A)** Schematic of mouse lever-press probabilistic reversal learning (PRL) task. Two levers extend on each trial; one delivers chocolate milk reward with 80% and the other 20% probability. Reward contingencies reverse after five consecutive responses on the high-probability lever. **(B)** Group-level three-state GLM-HMM fit to mouse behavior. Left, state transition matrix showing strong diagonal dominance indicating persistent latent states across trials. Right, state-specific GLM weights for bias, recent choice history, and recent rewarded-choice history. States were classified as *choice-persistence* (blue), *reward-achievement* (red), and *reward-learning* (green). **(C)** Comparison of *reward-learning* strategy weights estimated from the group model and subject-level individual fits, showing reliable recovery of the *reward-learning* strategy across animals. **(D)** Sequential GLM-HMM fits using sequentially increasing sets of training data (day 1 to day 12). Side biases dominate early behavior, whereas the three canonical latent states emerge by day 3 and remain stable thereafter. Reward sensitivity selectively increases in the *reward-learning* strategy across training. **(E)** Reward probability as a function of choice-streak length for each latent state. *Reward-achievement* and *reward-learning* strategies track task contingencies, peaking at the reversal criterion (five consecutive rewarded responses), whereas *choice-persistence* has persistently low reward probability across streak lengths. **(F)** Reward rate by latent state. *Reward-learning* and *reward-achievement* strategies yield higher reward rates than *choice-persistence*. **(G)** Reversal rate by latent state. Reversal frequency is highest in *reward-learning*, followed by *reward-achievement* and *choice-persistence*. **(H)** Reaction time (RT) by latent state. Responses are fastest during *reward-learning*. **(I)** Contingency policy space defined by stay–switch bias and contingency sensitivity index. *Reward-learning* uniquely combines strong feedback sensitivity with adaptive directional bias. **(J)** Simulation framework in which synthetic agents operate exclusively under each latent state using state-specific GLM weights as the generative policy. **(K)** Distribution of trials to reversal criterion in simulations. *Reward-learning* agents reach criterion fastest, *reward-achievement* agents are intermediate, and *choice-persistence* agents slowest. **(L)** Reward efficiency in simulated left) and empirical (right) data. Efficiency is highest for *reward-learning* strategies, intermediate for *reward-achievement* intermediate, and lowest for *choice-persistence*. Bars represent mean ± SEM; dots indicate individual mice. *p < 0.05, **p < 0.01, ***p< 0.001, ****p < 0.0001.

To identify underlying behavioral strategies, we fit a generalized linear model–hidden Markov model (GLM-HMM) to trial-by-trial choices to identify latent states, using recent choice and rewarded-choice history as predictors to allow latent states to capture distinct influences of reinforcement and action history on current decisions. Model comparison across varying state numbers using likelihood and information criteria supported a three-state solution (fig. S1). The corresponding transition matrix was strongly diagonally dominant (Fig. 1B), indicating that states persisted across multiple consecutive trials rather than fluctuating randomly from trial to trial, indicating that behavior reflects organized, stable strategies.

The three latent states were readily interpretable from their GLM weights (Fig. 1B). The first state strongly weighted recent rewarded choices and was named as a ‘*reward-learning*’ strategy. The second state showed weaker yet reliable reward dependence and was named as a ‘*reward-achievement*’ strategy. The third state was dominated by previous choice history with minimal reward sensitivity and was named as ‘*choice-persistence’*. Subject-level fits initialized from the group model reliably recovered the *reward-learning* strategy across animals (Fig. 1C), whereas the remaining states showed greater inter-individual variability (fig. S1). Individual transition matrices also retained strong self-transition probabilities, indicating sustained engagement of reward-related strategies across trials.

To examine how these strategies emerged across training, we sequentially refit the GLM-HMM while sequentially adding sessions across days. Early behavior was dominated by outcome-insensitive side biases, consistent with naïve exploration. By day three, all three canonical latent states became detectable and remained stable thereafter (Fig. 1D). Across training, the *reward-achievement* and *choice-persistence* strategies changed little, whereas the *reward-learning* strategy showed a marked increase in the previous rewarded trial regressor weight. Thus, experience did not simply increase response vigor; rather, learning selectively strengthened a latent strategy to prioritize recent reward outcomes in guiding future choice.

### Functional profiles distinguish latent behavioral strategies

We next asked whether the inferred latent states mapped onto differing profiles of observable task metrics. State-specific reward rates and feedback-adaptive metrics revealed marked functional differences across states (Table 1). Reward rate and lose–shift probability, defined as the probability of switching to the alternative option after an unrewarded outcome, were lowest for the *choice-persistence* strategy, consistent with a maladaptive perseverative policy. By contrast, lose-shift probability was substantially greater for the *reward-achievement* and *reward-learning* strategies indicating increased sensitivity to negative feedback and higher reward yield.

**Table 1.** Behavioral performance metrics vary by state in mice.

| Strategies | Reward rate | P(correct choice) | Win-stay | Lose-shift | CSI | Stay-switch bias |
| --- | --- | --- | --- | --- | --- | --- |
| Choice persistence | 0.349 | 0.249 | 0.662 | 0.232 | -0.106 | 0.43 |
| Reward achievement | 0.544 | 0.574 | 0.673 | 0.521 | 0.194 | 0.152 |
| Reward learning | 0.586 | 0.644 | 0.926 | 0.568 | 0.494 | 0.358 |

To further quantify reward efficiency, we calculated reward probability as a function of choice-streak length, defined as the number of consecutive selections of the same option (Fig. 1E). Under the *choice-persistence* strategy, reward probability remained uniformly low across streak lengths and gradually declined as streak length increased, reaching only ∼50% even at a streak length of one. Thus, animals persisted in repeating choices without effectively exploiting the rewarded option. In contrast, the *reward-achievement* and *reward-learning* strategies closely tracked task contingencies: reward probability increased across consecutive correct choices, peaked at the reversal criterion of five responses, and sharply declined when streak length exceeded this threshold. These profiles indicate adaptive maintenance of choice repetition within the beneficial range. Unlike reward-sensitive strategies, *choice-persistence* promoted perseveration beyond the useful extent of repetition, thereby reducing reward efficiency. Consistent with this interpretation, adding data from a subset of animals with one-sided choice availability increased the posterior occupancy and weight of the *choice-persistence* strategy (fig. S2), further validating this strategy as a perseverative policy.

Statistical comparisons confirmed that behavioral performance differed robustly across strategies. Reward rate varied by strategy [one-way ANOVA, F(2,59) = 107.73, p < 0.001] (Fig. 1F) and was lower during *choice-persistence* than *reward-achievement* or *reward-learning* (both p < 0.001) and the reward-sensitive strategies did not differ. Reversal frequency also depended strongly on strategy [F(2,59) = 61.90, p < 0.001] (Fig. 1G). *Choice-persistence* showed fewer reversals than *reward-achievement* and *reward-learning* strategies (both p < 0.001), and *reward-learning* exceeded *reward achievement* (p < 0.001). Reaction time similarly differed across states [F(2,59) = 23.41, p < 0.001] (Fig. 1H). Responses were faster during *reward-learning* than during *choice-persistence* or *reward-achievement* (both p < 0.001), whereas the latter two strategies did not differ significantly (p = 0.222).

To quantify how feedback history guided decision updating, we next derived two complementary metrics from win-stay (WS) and lose-shift (LS) probabilities. A Contingency Sensitivity Index [CSI = WS + LS − 1] measures the extent to which a strategy both exploits rewarded outcomes and updates behavior after non-reward. CSI was highest for the *reward-learning* strategy and lowest for *choice-persistence*, indicating maximal feedback sensitivity in the former and minimal adaptive updating in the latter. Because CSI alone cannot distinguish true feedback-guided responding from cases in which staying and switching occur at similar rates (WS ≈ LS), we additionally calculated a stay–switch bias (WS − LS) to capture directional tendencies independent of outcome. Mapping latent states within this WS–LS contingency policy space (Fig. 1I) showed that only the *reward-learning* strategy combined strong contingency sensitivity with a directional bias aligned to feedback history rather than symmetric or perseverative responding. Thus, *reward-learning* defined the most efficient strategy for simultaneously maximizing reward and adapting to reversals. Together, these analyses identify the *reward-learning* strategy as the most efficient behavioral policy, characterized by the highest reward rate, greatest reversal frequency, and fastest responding. In contrast, the *choice-persistence* strategy was characterized by inflexible responding and poorest overall performance.

### Simulations validate state-specific decision policies

Across trials, mice transition dynamically between latent states such that the behavioral consequences of any single strategy cannot be fully isolated in empirical data. We therefore simulated agents constrained to operate exclusively within each GLM-HMM state, using state-specific GLM weights to define the generative strategy (Fig. 1J). Simulated agents experienced the original trial structure and reward contingencies of the PRL task, permitting direct comparison between latent-state strategies in synthetic and real behavior.

State-restricted simulations closely reproduced the functional hierarchy observed *in vivo*. Across 6000 simulated trials, the *reward-learning* agent completed the greatest number of reversals (379), followed by *reward-achievement* (209) and *choice-persistence* (154), indicating superior adaptation to contingency changes under the *reward-learning* strategy. We next quantified the number of trials required to reach reversal criterion. The resulting trials-to-criterion distributions revealed a clear ordering, with *reward-learning* agents reaching criterion fastest, *reward-achievement* agents showing intermediate performance, and *choice-persistence* agents requiring the most trials (Fig. 1K; Kruskal–Wallis test, H = 114.69, P < 0.0001).

To quantify reward optimization, we computed a reward-efficiency score defined as accrued reward relative to the theoretical maximum obtainable under task contingencies. Reward efficiency showed the same hierarchy in empirical data and simulations (Fig. 1L). In mice, reward efficiency was highest during *reward-learning* (73.3%), intermediate during *reward-achievement* (68.0%), and lowest during *choice-persistence* (43.6%). Simulated agents closely recapitulated this structure (70.0%, 67.0%, and 47.6%, respectively).

Overall, simulation validated the interpretation of GLM-HMM states as meaningful, computationally distinct decision strategies. The close correspondence between simulated and observed behavior indicates that the latent states capture substantive strategies rather than simply clustering variability. In particular, the *reward-learning* strategy approximated the most adaptive strategy for simultaneously maximizing reward acquisition and efficient reversal performance.

**Table 2.** Simulated Agents Recapitulate State-Specific Mouse Behavioral Performance.

| Strategies | Simulated<br>Reward<br>rate | Real<br>Reward<br>rate | Simulated<br>P(correct<br>choice) | Real<br>P(correct<br>choice) | Reversal<br>numbers | Mean<br>trial to<br>criterion |
| --- | --- | --- | --- | --- | --- | --- |
| Choice persistence | 0.381 | 0.349 | 0.303 | 0.249 | 154 | 39 |
| Reward achievement | 0.536 | 0.544 | 0.564 | 0.574 | 209 | 28.7 |
| Reward learning | 0.560 | 0.586 | 0.594 | 0.644 | 379 | 15.8 |

### Humans employ shared and distinct latent decision strategies

We then asked if humans employ similar or distinct strategies in reward-based decision making. To test this, human participants (n = 26; 13 female) performed an analogous PRL task in which they selected between two abstract symbols presented on a computer screen using left or right button presses in a single session (Fig. 2A). One symbol was associated with an 80% probability of monetary gain and a 20% probability of loss, and another symbol carried the inverse contingencies. To increase task complexity while maintaining environmental volatility comparable to that experienced by mice, reversals were triggered probabilistically: following five correct choices within the previous six trials, each subsequent trial carried a 20% chance of contingency reversal.

**Fig. 2.**
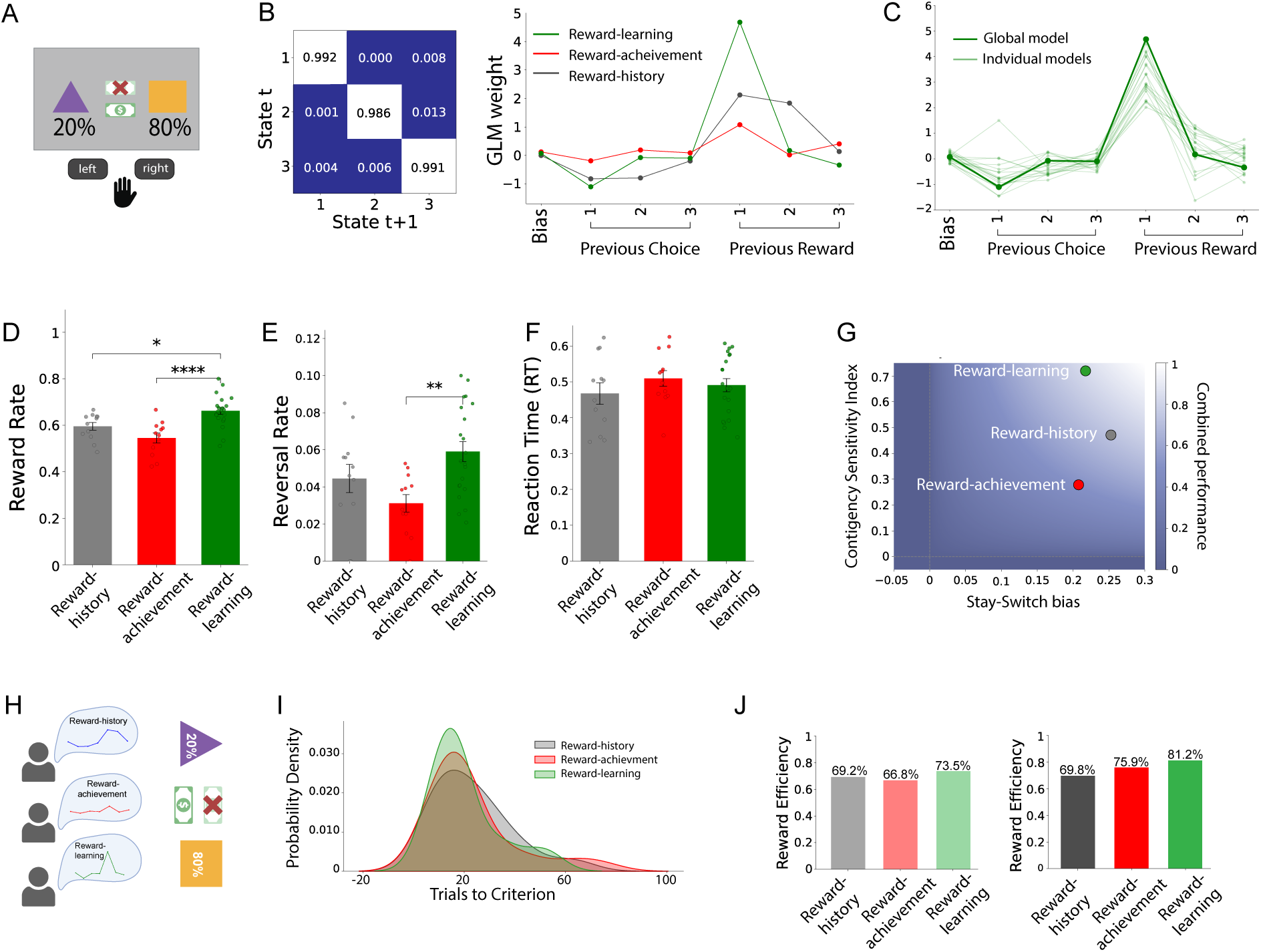
Humans engage shared and distinct latent decision strategies during probabilistic reversal learning. **(A)** Schematic of the human probabilistic reversal learning (PRL) task. Participants selected between two abstract symbols using left or right button presses. One symbol was associated with an 80% probability of monetary gain and 20% probability of loss and the other symbol carried inverse contingencies. After five correct choices within the previous six trials, each subsequent trial carried a 20% probability of contingency reversal. **(B)** Group-level three-state GLM-HMM fit to human behavior. Left, transition matrix showing strong diagonal dominance and persistent latent states across trials. Right, state-specific GLM weights for bias, recent choice history, and recent rewarded-choice history. States were classified as *reward-history* (gray), *reward-achievement* (red), and *reward-learning* (green). **(C)** Comparison of *reward-learning* strategy weights estimated from the group model and subject-level fits, showing reliable recovery of the *reward-learning* strategy across participants. **(D)** Reward rate by latent state. *Reward-learning* yielded higher reward rates than *reward-history* or *reward-achievement*. **(E)** Reversal rate by latent state. *Reward-learning* showed higher reversal frequency than *reward-achievement*, with *reward-history* intermediate. **(F)** Reaction time (RT) by latent state. No significant differences in response speed were observed across strategies. **(G)** Contingency policy space defined by stay–switch bias and contingency sensitivity index. *Reward-learning* combined strong feedback sensitivity with adaptive directional bias. **(H)** Simulation framework in which synthetic agents operated exclusively under each latent state using state-specific GLM weights as the generative policy. **(I)** Distribution of trials required to reach reversal criterion in simulations. *Reward-learning* agents reached criterion numerically fewer trials than *reward-history* and *reward-achievement* agents, although there was no significant difference. **(J)** Reward efficiency in simulated and empirical data. *Reward-learning* strategies showed the highest efficiency, followed by *reward-achievement* and *reward-history* strategies. Bars represent mean ± SEM; dots indicate individual participants. *p < 0.05, **p < 0.01, ***p < 0.001, ****p < 0.0001.

To compare cross-species learning performance, we computed a learning index quantifying the influence of prior reward on subsequent choice (see Methods) and confirmed that humans outperform mice (Fig. S3; t(36.9) = −6.20, p < 0.001, Hedges’ g = −1.68), indicating faster and more robust task acquisition. Because this metric is normalized by overall switching tendency, species differences could reflect enhanced reward sensitivity, altered persistence, or both. We therefore examined staying and switching separately after rewarded and unrewarded outcomes. Relative to mice, humans were more likely to repeat rewarded choices and more likely to switch after unrewarded outcomes (both p < 0.001), consistent with more reliable reinforcement-guided behavior rather than nonspecific exploration or disengagement. No significant sex differences were observed for any behavioral metric in either species (all p > 0.05) (fig. S3). Together, these findings suggest that superior human performance arises from more consistent use of outcome feedback.

We next asked whether humans and mice used the same latent strategies or species-specific policies. Applying the GLM-HMM to human trial-by-trial choices again supported a three-state solution (Fig. 2B; Fig. S4). Humans expressed latent states closely resembling the mouse *reward-achievement* and *reward-learning* strategies. However, instead of a *choice-persistence* strategy, humans exhibited a third state that strongly weighted the two most recent rewarded outcomes, which we termed *reward-history*. This difference likely reflects adaptation to task structure: whereas mice experienced deterministic reversals after five consecutive correct responses, humans learned under a probabilistic five-of-six criterion, giving utility to integration of short-term *reward-history* rather than simple persistence.

Despite this, the core reward-sensitive strategies were highly conserved. GLM weights for *reward-learning* and *reward-achievement* strategies were strongly correlated between mice and humans (*reward-learning*: r = 0.99, p < 0.0001; *reward-achievement*: r = 0.92, p = 0.003), indicating conserved internal decision strategies despite differences in overt behavior and task demands. As in mice, subject-level fits reliably recovered the *reward-learning* strategy across individual human participants (Fig. 2C), whereas the remaining strategies showed greater inter-individual variability (Fig. S5).

### Functional profiles distinguish latent strategies in humans

We next asked whether the latent states identified in humans corresponded to dissociable behavioral strategies. To address this, we compared reward rate, reversal rate, and RT across the three inferred states (Fig. 2, D to F). Reward rate differed significantly across strategies [one-way ANOVA, F(2,42) = 12.07, p < 0.001], and was higher under *reward-learning* than either *reward-history* (p = 0.025) or *reward-achievement* (p < 0.001) strategies, which did not differ from each other (p = 0.170). This indicates that *reward-learning* also associates with superior reward optimization in humans.

Behavioral flexibility, measured by reversal rate, differed significantly by strategy [F(2,42) = 5.67, p = 0.007] with higher reversal rates in the *reward-learning* strategy relative to *reward-achievement* (p = 0.005). However, reversal performance under the *reward-history* strategy did not differ from either *reward-achievement* (p = 0.345) or *reward-learning* (p = 0.206). Thus, the *reward-learning* strategy selectively associates with enhanced learning of the task rule. In contrast to mice, reaction time did not differ across human latent states [F(2,42) = 0.69, p = 0.507]. Accordingly, latent-state differences in humans were expressed primarily through reward maximization and adaptive switching rather than motor speed.

We next quantified feedback-guided decision updating using win-stay (WS) and lose-shift (LS) measures. As in mice, the Contingency Sensitivity Index [CSI = WS + LS − 1] was highest for the *reward-learning* strategy and lowest for *reward-history*, indicating that *reward-learning* most strongly integrates positive and negative feedback to guide future choices. Mapping strategies within the WS–LS contingency policy space (Fig. 2G) similarly showed only the *reward-learning* strategy combined high contingency sensitivity with a directional bias aligned to feedback history rather than symmetric responding.

Taken together, these findings demonstrate that the latent states recovered in humans reflect behaviorally meaningful decision strategies. Similar to observations in mice, the *reward-learning* strategy is the most efficient overall strategy, characterized by the highest reward rate and enhanced reversal performance.

**Table 3.** State-specific behavioral performance measures in humans.

| Strategies | Reward rate | P(correct choice) | Win-stay | Lose-shift | CSI | Stay-switch bias |
| --- | --- | --- | --- | --- | --- | --- |
| Reward history | 0.597 | 0.574 | 0.743 | 0.535 | 0.278 | 0.208 |
| Reward achievement | 0.607 | 0.679 | 0.862 | 0.609 | 0.471 | 0.253 |
| Reward learning | 0.649 | 0.749 | 0.968 | 0.751 | 0.719 | 0.217 |

### Simulations validate state-specific decision policies in humans

To isolate the behavioral relevance of each latent state, we again simulated agents constrained to operate exclusively under a single GLM-HMM state, using the corresponding state-specific GLM weights as the generative decision policy (Fig. 2H). State-restricted simulations reproduced the principal ordering observed in empirical behavior. Across 500 simulated trials, *reward-learning* and *reward-history* agents each completed 22 reversals, exceeding the *reward-achievement* agent (17 reversals), consistent with greater sensitivity to changing contingencies. Differences in number of trials required to reach reversal criterion were not statistically significant (Fig. 2I; Kruskal–Wallis test, H = 0.3813, P = 0.826), consistent with the more modest separation among latent strategies in human reversal performance.

To quantify reward optimization, we calculated reward efficiency as accrued reward relative to the theoretical maximum obtainable under task contingencies and reproduced the same ranking in simulated and empirical behavior (Fig. 2J). In human participants, reward efficiency was highest during *reward-learning* (81.2%), followed by *reward-achievement* (69.8%) and *reward-history* (66.6%). Simulated agents recapitulated this ordering (73.5%, 69.2%, and 66.6%, respectively), indicating that the inferred latent states correspond to computational differences in reward maximization. These simulations confirm that, as in mice, *reward-learning* approximates the most effective strategy for maximizing reward in humans. However, the relatively small differences between *reward-learning* and *reward-history* suggest that human performance may rely less on a single dominant policy and rather on flexible recruitment of multiple efficient strategies depending on task demands. More broadly, the close correspondence between simulated and observed behavior supports the interpretation of GLM-HMM latent states as distinct and interpretable decision-making strategies.

**Table 4.** Simulated Agents Broadly Recapitulate State-Specific Human Behavioral Performance.

| Strategies | Simulated<br>Reward<br>rate | Real<br>Reward<br>rate | Simulated<br>P(correct<br>choice) | Real<br>P(correct<br>choice) | Reversal<br>numbers | Mean trial<br>to criterion |
| --- | --- | --- | --- | --- | --- | --- |
| Reward<br>history | 0.554 | 0.597 | 0.618 | 0.574 | 22 | 22.6 |
| Reward<br>achievement | 0.534 | 0.607 | 0.608 | 0.679 | 17 | 23.1 |
| Reward<br>learning | 0.588 | 0.649 | 0.642 | 0.749 | 22 | 20.4 |

### Reward-learning strategy follows conserved logistic growth dynamics across species

To characterize the evolution of latent decision strategies, we examined posterior state probabilities across trials, averaged across subjects within each species (Fig. 3, A and B). In both mice and humans, the *reward-learning* strategy was initially expressed at low probability but increased progressively across trials, forming a characteristic warm-up trajectory. This rise emerged earlier in humans, becoming prominent near the beginning of the session, whereas in mice the transition occurred later, often towards the end of training. Comparable trajectories were also evident at the level of individual subjects (Fig. 3C,D), indicating that the population-level pattern was not driven by averaging alone.

**Fig. 3.**
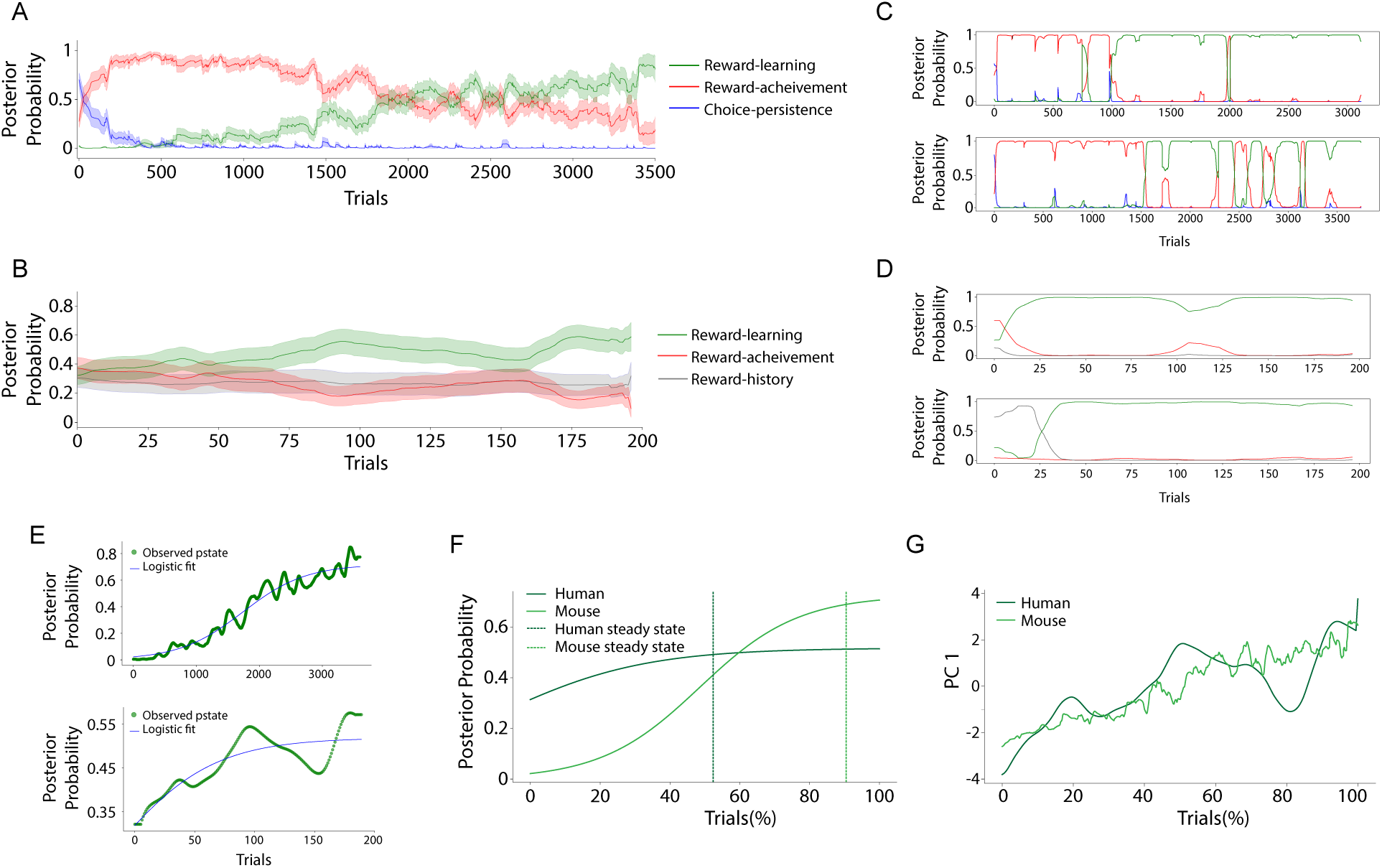
*Reward-learning* strategy emerges through conserved logistic growth dynamics across species. **(A)** Mean posterior probability trajectories across trials in mice for the three latent states identified by the GLM-HMM: *reward-learning* (green), *reward-achievement* (red), and *choice-persistence* (blue). Early behavior was dominated by *choice-persistence* and *reward-achievement*, whereas *reward-learning* gradually increased and dominated in later training. **(B)** Mean posterior probability trajectories across trials in humans for *reward-learning* (green), *reward-achievement* (red), and *reward-history* (gray). *Reward-learning* increased early in the session and increasingly surpassed the alternative strategies. **(C)** Representative posterior trajectories for individual mice illustrating subject-level emergence of *reward-learning* across trials. **(D)** Representative posterior trajectories for individual humans illustrating subject-level emergence of *reward-learning* within session. **(E)** Logistic growth model fits to *reward-learning* posterior probability over trials. Top, mice; bottom, humans. Smoothed empirical trajectories (green) are shown with fitted logistic curves (blue). **(F)** Direct comparison of species-level logistic growth models. Humans transitioned more rapidly into *reward-learning*, whereas mice showed slower growth but converged on a higher asymptotic probability of *reward-learning*. Dashed vertical lines indicate the estimated trial percentage at which steady state was reached for each species. **(G)** Projection of latent-state posterior trajectories onto the first principal component (PC1) across normalized trial progression. In both species, PC1 increased progressively over time and tracked *reward-learning* emergence, indicating a shared low-dimensional learning axis. Shaded regions in **(A)** and **(B)** represent SEM across subjects. Trial number in mice is normalized across sessions and shown as cumulative training progression; human trial number is normalized within session. Colors are consistent across panels.

Early behavior differed across species. In humans, initial performance was dominated by the *reward-achievement* strategy, which declined rapidly as *reward-learning* emerged, whereas the *reward-history* strategy remained comparatively stable across the session. In mice, early trials were dominated by the *choice-persistence* strategy, consistent with choice preservation. This tendency decayed rapidly and was followed by transient occupancy of the *reward-achievement* strategy before *reward-learning* gradually became dominant. Thus, although early strategy composition differed between species, both humans and mice ultimately converged toward increasing use of the *reward-learning* strategy.

The temporal organization of state usage also differed qualitatively. Mouse behaviour tended to be dominated by a single latent state at any given point, with gradual transitions toward increasingly reward-sensitive strategies. In contrast, humans showed greater co-occupation of multiple strategies, with substantial early overlap between *reward-achievement* and *reward-history* while *reward-learning* progressively increased to ultimately dominate. Despite these differences in strategy coordination, emergence of *reward-learning* was similar in both species. To quantify this transition, we modeled *reward-learning* posterior probability over trials using a dynamic logistic growth framework. Subject-wise reward-learning trajectories were first smoothed using a Kalman filter, averaged across subjects, and further denoised with a rolling mean. Logistic functions were then fit to the resulting group-level trajectories by nonlinear least squares (Fig. 3E). In humans, the logistic model provided a strong fit (RMSE = 0.0348), capturing the rapid early rise in *reward-learning* probability. Phase portrait analysis identified a stable steady-state value of 0.519, reached at approximately 52% of trials.

Applying the same framework to mice also yielded a good fit (RMSE = 0.0534). In mice, the steady-state *reward-learning* probability was higher (0.719) but was reached substantially later, at approximately 81% of trials. Comparison of fitted parameters showed that humans exhibited a markedly faster growth rate (0.024) than mice (0.0023), and a higher initial reward-learning occupancy (0.31 versus 0.0158). These findings indicate that humans engaged reward-guided strategies earlier and transitioned more rapidly away from naïve behavior, whereas mice required substantially more trials to overcome early *choice-persistence* before converging on *reward-learning* (Fig. 3F). No sex differences were detected in fitted growth dynamics in either species (Fig. S6).

To determine whether the emergence of *reward-learning* reflected a broader reorganization of strategy space, we next applied principal component analysis (PCA) to Kalman-smoothed posterior trajectories across all latent states. In both species, the first principal component (PC1) explained the largest amount of variance (humans: 82.3%; mice: 69.1%). PC1 increased progressively across trials and closely tracked *reward-learning* dynamics (Fig. 3G). Loadings on PC1 were positive for *reward-learning* and negative for all other strategies, indicating that this axis captures a dominant transition from early, less optimal policies toward more optimal reward-guided behavior. Although the specific alternative strategies differed across species, the alignment of PC1 with *reward-learning* was conserved.

Together, these findings identify *reward-learning* as an increasingly dominant strategy that governs behavior as subjects acquire the task. The similar logistic growth pattern across species supports *reward-learning* as a conserved computational policy whose emergence reflects gradual policy acquisition. Despite species-specific differences in early strategy composition, learning in both mice and humans unfolded along a common low-dimensional axis dominated by *reward-learning* dynamics.

### PRL engages conserved *reward-learning* strategy in mice across task modalities

Touchscreen operant chambers are increasingly used as a translational platform for assessing rodent cognition. To test whether touchscreen-based PRL captures the same latent decision strategies observed in lever-based PRL in mice and in computer-based human tasks, we analyzed behavior from mice trained in a touchscreen PRL task. Animals progressed through pre-training stages involving simple reward association, punishment of incorrect responses, and deterministic reversal learning before entering the probabilistic phase. In a total of seven sessions of PRL each up to 90 trials, mice (n=27, 15 female) selected between two visual stimuli under an 80/20 reward schedule with contingencies reversing after five consecutive correct choices (Fig. 4A).

**Fig. 4.**
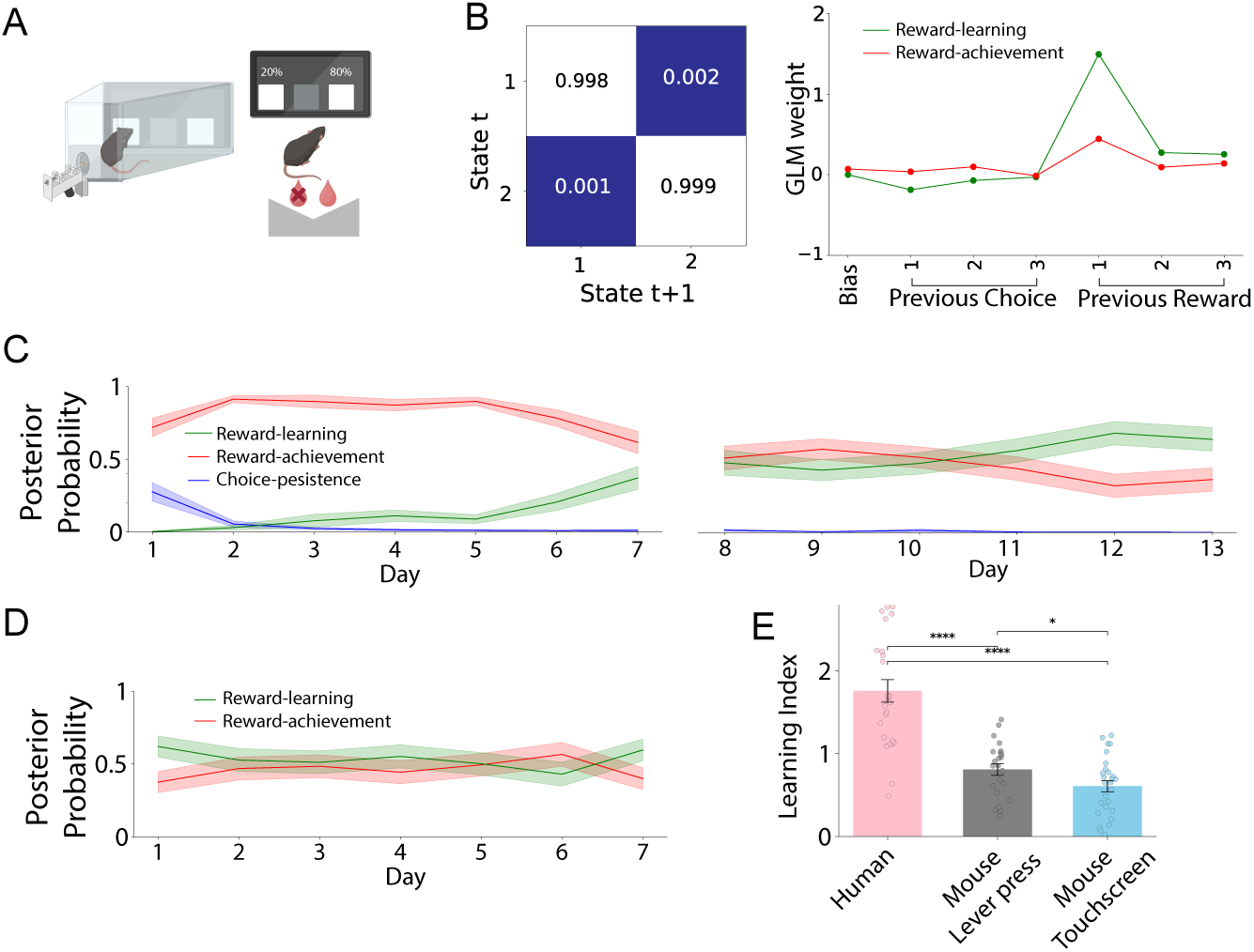
Touchscreen probabilistic reversal learning recovers conserved reward-sensitive strategies across task modalities. **(A)** Schematic of the mouse touchscreen probabilistic reversal learning (PRL) task. Mice selected between two visual stimuli presented on a touchscreen under an 80/20 reward schedule. Correct responses delivered reward, incorrect responses produced negative feedback, and contingencies reversed after five consecutive correct choices. **(B)** Group-level two-state GLM-HMM fit to touchscreen PRL behavior. Left, transition matrix showing strong diagonal dominance and stable latent states across trials. Right, state-specific GLM weights for bias, recent choice history, and recent rewarded-choice history. The recovered states corresponded to *reward-learning* (green) and *reward-achievement* (red). **(C)** Posterior state probabilities across sessions. During early training, *choice-persistence* (blue) declined rapidly, *reward-achievement* (red) dominated initial sessions, and *reward-learning* (green) progressively increased across days. Panels show days 1–7 (left) and 8–13 (right). **(D)** Posterior state probabilities across sessions for touchscreen PRL mice. *Reward-learning* and *reward-achievement* strategies stably co-express across sessions. **(E)** Learning index comparison across humans, lever-press PRL mice, and touchscreen PRL mice. Learning index is modestly higher in lever press than touchscreen mice, and both are lower than humans. Bars represent mean ± SEM; dots indicate individual subjects. Colors are consistent across panels: *reward-learning* (green), *reward-achievement* (red), and *choice-persistence* (blue). *p < 0.05, **p < 0.01, ***p< 0.001, ****p< 0.0001.

We next fit the same GLM-HMM framework used for lever-based mouse and human data with identical behavioral covariates. Model comparison supported a two-state solution in touchscreen PRL (fig. S7). These two latent states closely matched the previously identified *reward-learning* and *reward-achievement* strategies and exhibited strong state persistence (Fig. 4B), indicating that touchscreen behavior preserved the core reward-sensitive behavioral strategies observed across species and task formats. In contrast to lever-based PRL, a distinct *choice-persistence* strategy was not identified. *Choice-persistence* was prominent in early training in lever-based PRL. Potentially this strategy may be reduced by the extended pretraining specific to touchscreen tasks, or alternatively by stimulus-response mapping, or modality-dependent task demands.

To compare strategy dynamics across mouse task variants, we then examined posterior state probabilities over training across tasks. Restricting to the first seven sessions of lever-based PRL to match touchscreen duration, *reward-learning* occupancy gradually increased while *reward-achievement* declined, and *choice-persistence* rapidly diminished (Fig. 4C). In contrast, touchscreen mice showed relatively stable co-expression of *reward-learning* and *reward-achievement* strategies that was maintained across training (Fig. 4D). Thus, although touchscreen PRL recovered the same reward-sensitive latent strategies the temporal evolution was distinct. We next compared learning performance across species and task modalities using the learning index. Lever-trained mice showed modestly higher learning indices than touchscreen-trained mice [t(47.4) = 2.06, p = 0.045, Hedges’ g = 0.57]. As expected, humans substantially outperformed both touchscreen mice [t(36.7) = 7.54, p = 5.76 × 10⁻⁹, Hedges’ g = 2.06] and lever-trained mice [t(36.9) = 6.20, p = 3.47 × 10⁻⁷,Hedges’ g = 1.68] (Fig. 4E). These findings indicate that lever-press and touchscreen PRL tasks produce broadly comparable levels of learning and across tasks, mice are substantially less efficient learners than humans. Despite deterministic pretraining and fewer trials per session in the touchscreen paradigm, touchscreen PRL nonetheless recovered reward-sensitive latent strategies closely aligned with those identified in lever-based mouse PRL and human PRL. The relatively stable balance of *reward-learning* and *reward-achievement* in the touchscreen task may reflect prior deterministic pretraining, reduced opportunity for strategy evolution across limited sessions, or apparatus-specific action costs, since animals must leave the response location to retrieve reward, making reward collection potentially more costly than changing choices. Overall, touchscreen PRL supports the cross-task robustness of reward-learning strategies while also suggesting that their temporal expression may depend on task history and motor demands.

## DISCUSSION

Here, we used probabilistic reversal learning and latent variable models (GLM-HMMs) to decompose trial-by-trial behavior to identify latent decision-making strategies across mice and humans. A *reward-learning* strategy emerged as the most optimal strategy, showing the highest reward efficiency, strongest sensitivity to changing contingencies, and greatest ability to adapt following reversals. This strategy was broadly conserved in both species, although with differing timescales, suggesting that reward learning represents a shared computational policy for acquiring and updating task rules in dynamic environments. Thus, PRL performance reflects both a conserved *reward-learning* strategy and species- or task-specific dynamics to converge toward a reward-guided policy.

These findings support a state-based view of reversal learning, in which behavior is better understood as transitions between latent decision policies rather than the output of a single fixed learning process. Performance metrics such as accuracy, reward rate, or switching probability offer limited insight into the strategies that generate choices (*12–14*) and similar performance can arise from distinct underlying processes (*15, 16*). Consistent with prior evidence that rodents use multiple behavioral modes during reversal learning (*3, 16, 17*), we found that both mice and humans transitioned between distinct latent states while converging on a conserved *reward-learning* strategy. Importantly, the timing, persistence, and frequency of state occupancy differed across individuals indicating that state-based modeling can capture individual variability in strategy deployment while revealing conserved computational policies that may be obscured by aggregate performance measures. GLM-HMM balances flexibility and interpretability by estimating discrete latent states defined by weights on observable covariates, allowing us to distinguish cases in which similar actions reflected different underlying policies. For example, repeating a choice could reflect adaptive exploitation after reward, perseverative choice stickiness, or integration of recent reward history, depending on the latent state. The use of common covariates across mouse and human tasks enabled direct comparison of latent strategy across species and task modalities. Although our model does not assign states to explicit computational agents, the resulting structure is compatible with the broader view advanced by mixture-of-agents HMMs (MoA-HMM), in which behavior reflects transitions among competing decision policies rather than noisy deviations from a single learning algorithm (*18*).

The conserved emergence of the *reward-learning* strategy suggests that optimal PRL performance depends not only on maximizing reward, but also on optimizing the strategy to obtain reward. PRL requires subjects to first identify and exploit the currently rewarded option and then second, to detect when the contingency has reversed and update behavior accordingly. A policy that captures only the first component can support high reward acquisition during stable periods but is inefficient when flexible adaptation is required by reversals. Thus, reward maximization and strategy optimization are not equivalent: subjects may obtain many rewards while still using a policy that is slower, more perseverative, or less aligned with the full task structure.

This distinction was most evident in mice. Early in training, mice expressed a *reward-achievement* strategy that supported reward acquisition by maintaining choices on the currently advantageous lever. This strategy likely captures the contingency component of the task: after rewarded choices, staying with the same option increases the probability of obtaining additional reward. However, this policy does not fully capture the reversal component. Once the reversal criterion is reached, persistence on the previously rewarded lever becomes inefficient and requires renewed exploration before the animal identifies the new high-probability option. In contrast, the *reward-learning* strategy incorporated both reward-guided repetition and sensitivity to reversing contingencies. Although reward rate increased only modestly during reward-learning occupancy, reversal tracking improved and reaction times decreased, indicating a more efficient, task-aligned mode of performance. In this sense, reward learning optimized the process by which reward was obtained, rather than simply increasing reward rate.

A similar principle was observed in humans, although auxiliary strategies differed across species. Humans showed less exclusive dominance of a single strategy and maintained greater access to multiple strategies across the task. The human-specific *reward-history* strategy performed comparably, and in some measures slightly better than the *reward-learning* strategy, suggesting that integrating reward information across multiple previous trials can be useful under probabilistic uncertainty. This may be especially relevant because the human task used a five-of-six reversal rule, in which a single unrewarded outcome does not necessarily indicate a reversal. However, maintaining reward information across a longer recent history may impose greater working-memory or computational demands than relying primarily on the most recent rewarded choice (*19*). *Reward-history* may therefore serve as an auxiliary strategy that increases robustness under uncertainty, whereas *reward-learning* provides a simpler, more general policy for aligning behavior with the core task structure. Consistent with this interpretation, human reaction times did not differ strongly across states, suggesting that state-dependent differences in humans were expressed primarily through reward maximization and reversal adaptation rather than motor response speed.

These findings suggest that *reward-learning* represents an optimized strategy for solving PRL across species. This conservation indicates that *reward-learning* is a robust and translatable behavioral strategy, preserved despite species differences in cognitive capacity, motor output, and task modality. These properties make *reward-learning* a strong candidate for future studies testing how stress, affective symptoms, or neural circuit manipulations alter reward-guided behavioral adaptation (*20, 21*).

A key insight from the cross-species analysis is that conservation of a strategy does not require conservation of timescale. Humans expressed *reward-learning* rapidly within one session, whereas mice showed across-session evolution of strategy structure. In mice, analyzing expanding session windows provided several converging observations supporting this process. First, modeling early sessions alone revealed behavior was largely captured by naïve side *bias*, consistent with initial responding driven by pre-existing choice preference than task-derived information. Second, adding additional sessions increased organization of state-specific weights around choice and reward history, particularly in the *reward-learning* state. Third, *reward-learning* posterior probabilities progressively increased, indicating that this strategy was not only more identifiable in model weights but also more frequently deployed as training progressed. This across-session dynamic also reveals an important cross-species distinction in the timescale of strategy formation. This may reflect species-specific cognitive capacity, task comprehension, prior experience, or the advantage of explicit instruction in humans (*22*). More broadly, these findings suggest that learning paradigms should be analyzed not only by pooling trials across sessions, but by tracking how accumulated experience changes how trial-level information shapes individual performance and how latent strategies reorganize across experience.

Several limitations should be considered. While prior work shows that males and females can achieve similar overall performance while relying on different explore–exploit strategies or responding differently to negative feedback (*23*), we did not observe significant sex differences in learning indices, choice-consistency measures, or logistic growth dynamics in either species. While we show that the emergence of *reward-learning* is broadly conserved across males and females, subtler sex-dependent differences in motivational state, trial initiation, or neural implementation might be revealed with larger samples. Further, although the mouse and human PRL tasks were comparable, they were not identical. Mice performed multiple lever-press or touchscreen sessions with criterion-based reversals, whereas humans completed a single computerized session with a probabilistic five-of-six reversal rule. These differences likely contributed to species-specific auxiliary states and to the different timescales of reward-learning emergence. In addition, the touchscreen dataset included extensive pretraining and fewer trials per session, which may have limited detection of slower learning dynamics or stickiness-related states. The root cause of timescale differences remains an open question that future parametric studies using closely matched paradigms, multi-session human PRL, or instruction-free human tasks could help resolve. Finally, because GLM-HMM states are inferred from behavior, they should be interpreted as latent behavioral strategies rather than direct evidence for specific cognitive or neural mechanisms. Future studies combining state inference with trial-level neural recordings and circuit manipulations can probe whether *reward-learning* and auxiliary strategies correspond to distinct neural activity patterns, whether state transitions are preceded by changes in circuit dynamics, and whether perturbing specific circuits can shift behavior toward or away from reward-guided control.

Here, we show that PRL behavior can be decomposed into latent decision strategies that differ in their reliance on choice history, reward history, and bias. Across mice and humans, *reward-learning* emerged as a conserved and optimized strategy for adapting to changing reward contingencies, while species and task differences were reflected primarily in the timescale and auxiliary strategies through which this policy was expressed. This framework defines a robust translational foundation to uncover how reward-guided behavioral adaptation is shaped by learning history, life experience, affective state, and the underlying neural circuit mechanisms.

## MATERIALS AND METHODS

### Mouse lever-press PRL

Twenty-three C57BL/6J mice (11 females) aged 7 weeks at the start of the experiment were housed under standard laboratory conditions on a 12 h light/dark cycle (lights on at 07:00), with ambient temperature maintained between 22–25 °C and relative humidity at approximately 50%. Mice were group-housed in same-sex cages of three to four with ad libitum food and water. All experimental procedures were approved by the McGill Animal Care Committee and complied with institutional and national guidelines for animal research.

### Lever-press training

Mice underwent a multistage lever-press training protocol consisting of three sequential stages, with each daily session lasting 30 minutes. Prior to behavioral training, mice were food restricted to 85–90% of their free-feeding body weight. In the first stage, animals were trained to associate lever pressing with reward delivery. At the start of each trial, both levers were extended and the cue lights above each lever were illuminated. Animals were given up to 60 s to press with a press on either lever resulting in lever retraction, immediate delivery of a 30 µL chocolate milk reward, and presentation of a 3 s auditory cue (2 kHz pure tone or white noise). If no response was made within the response window, the trial was recorded as an omission. All trials were followed by a 10 s intertrial interval (ITI). After completing one session with more than 25 responses, animals advanced to the next training stage. During the second stage, the trial structure remained identical, but reward delivery was probabilistic: presses on either lever resulted in reward with 50% probability. On rewarded trials, chocolate milk delivery was paired with a 3 s auditory cue, and on unrewarded trials no reward was delivered and a distinct auditory cue was presented. Cue identity (pure tone vs white noise) was counterbalanced across animals. Animals progressed to the third stage after completing two consecutive sessions with more than 40 responses. The third stage was identical to the second stage, except that the response window was reduced to 10 s. Following two consecutive sessions with more than 100 responses, animals advanced to the two-armed bandit task.

### Two-armed bandit task

The two-armed bandit task was conducted over six consecutive days, with each session lasting 1 h. At the beginning of each trial, both levers were extended, and the associated cue lights were illuminated. Pressing a lever resulted in immediate lever retraction and delivery of the trial outcome. One lever was associated with an 80% probability of reward, whereas the other was rewarded on 20% of trials. Rewarded trials consisted of a 30 µL chocolate milk reward paired with a 3 s auditory cue, while unrewarded trials were signaled by a distinct 3 s auditory cue. Cue–outcome associations were counterbalanced across animals (maintained from pre-training). If no press was made within10 s, an omission was registered. Each trial was followed by a 10 s ITI. To promote continuous learning and frequent contingency updating, reward probabilities were reversed between levers after five consecutive responses on the currently high-probability lever.

### Human experiment

Twenty-six healthy adults (13 females) participated in the human experiment. All participants provided written informed consent and basic demographic information prior to participation. Experimental procedures were approved by Regional Ethics Review Board in Linköping, Sweden Dnr 2016/497-31and were conducted in accordance with the Declaration of Helsinki.

Participants completed a probabilistic reversal learning (PRL) task in which two abstract symbols were presented on each trial. On every trial, participants selected one symbol using a button press and were instructed to maximize their monetary earnings by choosing the option they believed to be currently most advantageous. One symbol was associated with an 80% probability of monetary gain and a 20% probability of monetary loss, whereas the other symbol had the complementary probabilities. The spatial positions of the symbols (left vs right) were randomized across trials. Reversal of reward contingencies was performance dependent. After participants achieved at least five correct choices within the previous six trials, the task entered a reversal-eligible phase, during which the reward contingencies could switch between symbols with a probability of 0.2 on each subsequent trial. Each trial consisted of a stimulus presentation period (1.5 s), during which participants were required to select, followed by outcome feedback (1 s). Intertrial intervals were jittered and drawn from an exponential distribution ranging from 1 to 6.5 s. If no response was made within the stimulus window, the trial was recorded as an omission and the message “Too slow!” was displayed. Upon response, a visual frame highlighted the selected symbol, and feedback was presented as either a coin image accompanied by the message “You won!” or a crossed-out coin with the message “You lost!”.

### Mouse touch screen PRL

Raw behavioural data from the touchscreen-based PRL task were obtained online through the open MouseBytes touchscreen database (*24–26*). Twenty-seven C57BL/6J mice (15 females, backcrossed onto the C57BL/6J background for more than ten generations), aged 7 weeks at the start of the experiment were tested in a touchscreen-based probabilistic reversal learning (PRL) paradigm adapted from the BrainsCAN Rodent Cognition Core standard operating procedure. Behavioral testing was conducted in automated touchscreen chambers (Lafayette Instrument, Lafayette, IN, USA) equipped with two response windows positioned on the far left and far right sides of the screen and controlled using ABET II software. A strawberry milkshake reward was delivered through a reward magazine accompanied by illumination of the tray light and an auditory tone.

Prior to behavioral training, mice were food restricted to 85–90% of their free-feeding body weight and habituated to the strawberry milkshake reward (Neilson Dairy, Mississauga, Ontario) in their home cages for at least 3 days. Animals then underwent a multistage touchscreen training procedure consisting of habituation, initial touch, must-touch, punish-incorrect, and deterministic reversal learning stages. During habituation, mice acclimated to the chambers and learned reward collection from the illuminated food tray. During initial touch and must-touch stages, mice learned to associate touchscreen responses with reward delivery. In the punish-incorrect stage, responses to the blank stimulus location resulted in a timeout period without reward, training mice to discriminate between response options.

Following pretraining, mice underwent deterministic reversal learning to familiarize them with within-session contingency reversals. Two identical white square stimuli were simultaneously presented in the left and right response windows, with one designated as the rewarded stimulus (S+) and the other as the unrewarded stimulus (S−). Responses to the S+ stimulus were always rewarded, whereas responses to the S− stimulus were never rewarded. After five consecutive responses to the rewarded stimulus, reward contingencies reversed such that the previously rewarded side became unrewarded and vice versa.

The PRL test proper used the same task structure but introduced probabilistic feedback contingencies. At the start of each trial, two white square stimuli were presented simultaneously in the left and right response windows. One side was designated as optimal (S+) and the other as suboptimal (S−). Selection of the optimal stimulus resulted in reward delivery on 80% of trials and omission of reward on 20% of trials, whereas selection of the suboptimal stimulus resulted in reward omission on 80% of trials and reward delivery on 20% of trials. Stimuli remained on the screen until a response was made. Correct responses were followed by reward delivery, magazine illumination, and an auditory tone, whereas incorrect or unrewarded responses resulted in illumination of the magazine light without reward delivery. Mice were required to enter the reward magazine before initiation of the inter-trial interval. After five consecutive responses to the optimal stimulus, contingencies reversed within the same session. Sessions lasted a maximum of 60 min and consisted of up to 90 trials.

### GLM-HMM Modeling of Latent Decision Strategies

To identify latent decision-making strategies underlying trial-by-trial choice behavior, we applied a Hidden Markov Model with Bernoulli Generalized Linear Model observations (*27*). This framework models behavior as transitions between discrete latent states, each associated with a distinct decision policy that maps recent trial history onto choice probabilities. Each latent state was characterized by a row of a state transition matrix and a corresponding vector of GLM weights, which determined the probability of selecting the right versus left option on each trial. The observation model included a set of covariates capturing recent behavioral history: (1) a constant bias term, (2–4) indicators encoding the subject’s choices on the three preceding trials, and (5–7) indicators encoding rewarded choices over the previous three trials.

We first fit GLM-HMMs to group-level behavioral data by concatenating trials across all subjects within each dataset. Model parameters were estimated using the Expectation– Maximization algorithm, and posterior state probabilities were inferred for each trial. The number of latent states was determined using a combination of cross-validated log-likelihood, Akaike Information Criterion (AIC), and Bayesian Information Criterion (BIC), with a three-state model providing the optimal balance between model fit and interpretability. Following group-level model estimation, the inferred transition matrix and GLM weights were used to initialize individual-level GLM-HMMs for each subject. This procedure enabled assessment of state engagement at the individual level while maintaining consistent initialization across subjects. Individual models converged reliably and exhibited state structures comparable to those observed at the group level. The same modeling and fitting procedures were applied independently to both the human and mouse datasets. Because mice completed the task across multiple sessions, GLM-HMM weights from the best-fitting model were additionally visualized as a function of cumulative training by progressively incorporating sessions over time, allowing assessment of how inferred decision policies evolved with experience.

### Behavioral performance metrics and statistical analysis

To characterize the functional profile of each latent behavioral strategy, we quantified reward rate, reversal rate, and reaction time (RT) separately for trials assigned to each GLM-HMM state. Trial-wise <u>latent state assignment</u> was determined using the maximum posterior probability inferred by the GLM-HMM. <u>Reward rate</u> was calculated as the proportion of rewarded trials within each latent state. <u>Reversal rate</u> was defined as the number of successful contingency reversals normalized by the total number of trials assigned to the corresponding state. For mice, reversals occurred after five consecutive correct responses on the high-probability option. For humans, reversals were probabilistic and occurred with 20% probability after five correct choices within the previous six trials. <u>Reaction time</u> was calculated as the latency between stimulus presentation and behavioral response (lever press in mice or button press in humans) and averaged within each state.

Differences in reward rate, reversal rate, and reaction time across latent states were assessed separately for mice and humans using one-way analysis of variance (ANOVA). When significant main effects were detected, pairwise comparisons were performed using Tukey’s honestly significant difference (HSD) post-hoc tests. For comparisons between species or sexes, independent-samples Welch’s t tests were used to account for unequal variance where appropriate. Effect sizes were quantified using Hedges’ g. Statistical significance was defined as p < 0.05. All analyses were performed in Python using *SciPy* and *statsmodels*.

### Policy-dependent reward rates and feedback-guided adaptation metrics

To quantify how latent decision strategies relate to task performance, we computed state-resolved reward rates and feedback-guided adaptation metrics using trial-wise posterior state probabilities from the GLM-HMM. On each trial *t*, choices were coded as *c_t_*(left/right; lever press) and outcomes were binarized as reward delivered versus not delivered, *r_t_* ∈ {0,1}, by recoding reward delivery as *r_t_* = I(reward_delivered*_t_* ≠ 0). The GLM-HMM provided posterior probabilities *w_t_*_,*k*_ = *p*(*z_t_* = *k* ∣ data)for each latent state *k*. All metrics were estimated using posterior-weighted (“soft”) assignments such that each trial contributed fractionally to each state in proportion to *w_t_*_,*k*_.

### Win–stay, lose–shift, and feedback sensitivity

To quantify immediate feedback-guided updating, we computed win–stay (WS) and lose– shift (LS) probabilities for each state *k*. The next-trial choice *c_t_*_+1_was defined by shifting the choice vector forward by one trial, and the final trial of each sequence was excluded. Trial-wise indicators were defined as win (I(*r_t_* = 1)), loss (I(*r_t_* = 0)), stay (I(*c_t_*_+1_ = *c_t_*)), and shift (I(*c_t_*_+1_ ≠ *c_t_*)). WS and LS were then computed using posterior-weighted counts:

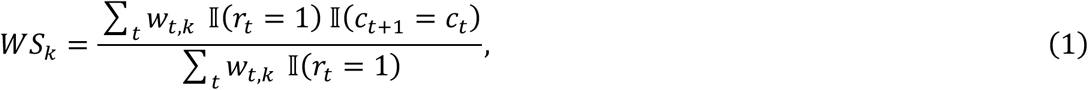

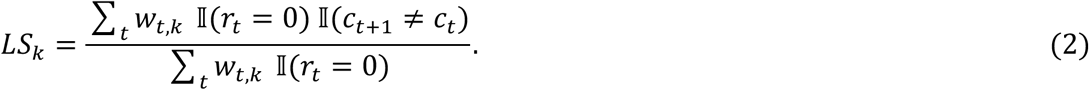

From these measures, we derived two complementary metrics. A Contingency Sensitivity Index (CSI) quantified the degree to which a policy both exploits reward and adapts following non-reward:

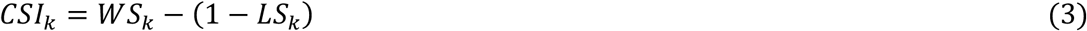

This index equals zero under chance-like, non-contingent responding (e.g., *WS* ≈ *LS* ≈ 0.5), increases when both reward exploitation and adaptive switching are present, and becomes negative when both are reduced.

To capture feedback-independent directional tendencies, we additionally computed a stay– switch bias:

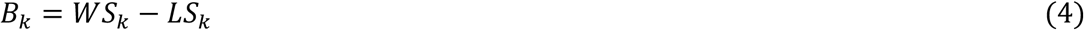

where positive values indicate a bias toward staying and values near zero reflect symmetric stay–switch behavior. For visualization in the WS–LS contingency policy space, each state was represented as a point (*LS_k_*, *WS_k_*), where CSI corresponds to movement along the diagonal and *B_k_* reflects displacement relative to the line *WS* = *LS*.

### Choice-streak–dependent reward probability

To assess how latent strategies interact with task structure, we quantified reward probability as a function of choice-streak length. Choice streak length *s_t_* was defined as the number of consecutive trials up to trial *t*on which the same option was selected:

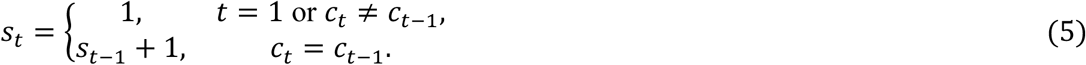

For each state *k*and streak length *s*, posterior-weighted reward probability was computed as:

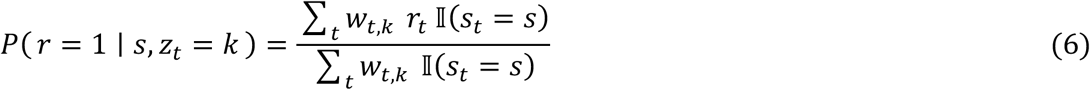

### Simulation of state-specific decision policies

Because empirical human and mouse behavior reflects dynamic switching between multiple latent strategies, the behavioral signatures of individual GLM-HMM states cannot be fully isolated from real data alone. To characterize the decision policy associated with each latent state in isolation, we simulated agents operating exclusively under each inferred GLM-HMM state, using the state-specific GLM weights as generative decision rules. This approach enabled direct comparison between empirical behavior and the behavior expected from each latent policy in its pure form.

Simulated agents were exposed to the same probabilistic reversal learning (PRL) task structure used in the empirical experiments. On each trial, agents selected between two options (left/right), one of which was designated as the currently correct option. Correct choices were rewarded with probability *P*(reward ∣ correct) = 0.8, whereas incorrect choices were rewarded with probability 0.2. A reversal occurred when the agent achieved a criterion of five consecutive correct choices. Upon criterion attainment, the identity of the correct option was reversed and the counter for consecutive correct responses was reset. Simulations were run for long sequences (mice: 6,000 trials; humans: 500 trials) to ensure stable estimation of reversal statistics and reward efficiency.

For each latent state *k*, agent behavior was generated using the GLM parameters inferred for that state from empirical data. On each trial *t*, the probability of choosing the left option according to the GLM was computed as

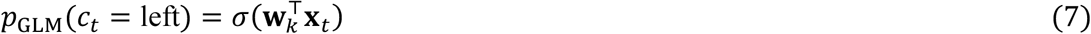

where *σ*(⋅)denotes the logistic sigmoid function, **w***_k_* are the GLM weights for state *k*, and **x***_t_* is a feature vector comprising a constant term, the previous three choices, and the previous three signed reward outcomes. Signed rewards were defined as the chosen action on rewarded trials and zero otherwise, consistent with the empirical GLM-HMM formulation.

To assess robustness of state-specific policies with respect to feedback-based heuristics, we additionally defined a win–stay/lose–shift (WSLS) policy using empirically estimated WS and LS parameters for each state. The WSLS probability of choosing left was defined conditionally on the previous choice and reward outcome. The final choice probability was computed as a convex combination:

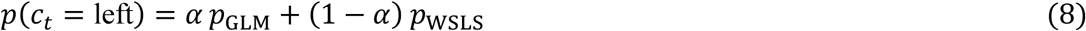

where *α* ∈ [0,1]controls the contribution of the GLM-based policy. Unless otherwise stated, simulations used *α* = 1, corresponding to purely GLM-driven behavior; alternative values produced qualitatively similar results. Choices were sampled stochastically on each trial from the resulting Bernoulli distribution.

### Performance metrics and statistical analysis

For each simulated agent, we quantified reward rate (fraction of trials with reward), accuracy (fraction of trials in which the correct option was selected), number of reversals (number of times the reversal criterion was reached), and trials to criterion (number of trials elapsed between reversals until five consecutive correct choices were achieved). Trials-to-criterion distributions were computed separately for each state-specific agent.

Differences in trials-to-criterion distributions across simulated agents were assessed using a nonparametric Kruskal–Wallis test, chosen due to non-normality and unequal variances across states. Statistical analyses were performed separately for mouse and human simulations.

To quantify how efficiently each policy extracted reward relative to the task maximum, we computed a reward efficiency score by normalizing each agent’s reward rate by the maximum achievable reward probability in the task (0.8). This metric reflects the proportion of optimal reward obtained by each policy and enables direct comparison between simulated and empirical performance.

### Dynamic modeling of latent strategy evolution

#### Posterior state trajectories

Latent state probabilities were obtained from the GLM-HMM using the Viterbi algorithm to infer trial-wise state sequences. For each trial, posterior probabilities over states were computed and used to characterize the temporal evolution of strategy engagement. In humans, posterior trajectories were derived from a single experimental session per subject. In mice, posterior trajectories were concatenated across all behavioral sessions for each subject to capture learning dynamics across extended training.

To visualize group-level dynamics, posterior state probabilities were first aligned by trial index within each subject and then averaged across subjects separately for humans and mice. This procedure yielded population-level trajectories describing how the probability of occupying each latent state evolved across trials.

#### Kalman smoothing of posterior trajectories

To reduce trial-wise noise while preserving temporal structure, subject-level posterior probability trajectories for each state were smoothed using a linear Gaussian state-space model implemented via a Kalman smoother. For each state and subject, posterior probabilities were modeled as noisy observations of an underlying latent process evolving according to:

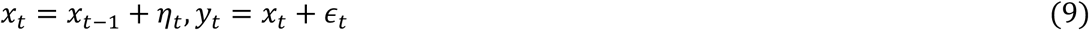

where *x_t_* denotes the latent smoothed probability, *y_t_* the observed posterior probability, and *η_t_* and *ε_t_* are zero-mean Gaussian process and observation noise, respectively. Transition and observation matrices were set to identity, with fixed process and observation noise parameters. Kalman smoothing was applied independently to each subject and each state.

Following smoothing, state probabilities were renormalized on each trial to ensure that probabilities summed to one across states. Smoothed trajectories were then averaged across subjects to obtain group-level state probability trajectories.

#### Logistic growth modeling of reward-learning emergence

To quantitatively capture the emergence of the reward-learning strategy over time, we modeled the group-averaged, Kalman-smoothed posterior probability of the reward-learning state using a logistic growth model. Prior to model fitting, a rolling mean was applied to the group-level trajectory to further reduce high-frequency variability.

The logistic growth model was defined as:

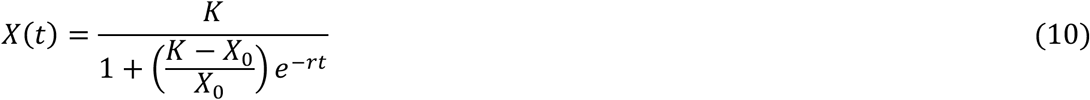

where *X*(*t*)is the posterior probability of the reward-learning state at trial *t*, *X*_0_ is the initial value, *K*is the asymptotic steady-state value, and *r* is the growth rate. Model parameters were estimated using nonlinear least-squares fitting. Model performance was assessed using root mean square error (RMSE).

To identify steady-state behavior, we examined the derivative of the fitted logistic function with respect to time. Fixed points were identified by solving 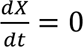, yielding an unstable fixed point at zero and a stable fixed point at *X* = *K*. The trial at which the fitted trajectory entered a neighborhood of the steady state was used to estimate the point of stabilization.

#### Low-dimensional analysis of strategy reorganization

To assess whether the emergence of the reward-learning state reflected a broader reorganization of strategy space, we applied principal component analysis (PCA) to the Kalman-smoothed posterior state probabilities. PCA was performed separately for humans and mice on group-averaged trajectories across all states. Prior to PCA, state probability trajectories were z-scored across trials to ensure comparable scaling.

The first principal component (PC1) was interpreted as a dominant learning axis, capturing the primary direction of variance in strategy usage over time. State loadings on PC1 were examined to determine which strategies contributed positively or negatively to this axis. Trial-wise projections onto PC1 were then used to visualize the temporal evolution of behavior in the reduced-dimensional space.

To facilitate cross-species comparison, PC1 trajectories were plotted as a function of normalized trial progression (trial index divided by total number of trials). This allowed direct comparison of learning dynamics between humans and mice despite differences in session length.

#### Behavioral Analysis: Reward Sensitivity and Learning Index

To characterize how reward feedback shapes decision updating in humans and mice, we quantified outcome-dependent switching behavior using a normalized learning index. For each subject, we computed the conditional probability of switching choices following rewarded and unrewarded trials, and defined the learning index as:

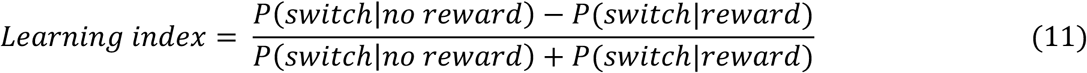

This metric captures sensitivity to feedback by measuring the extent to which negative outcomes promote switching relative to positive outcomes, while controlling for individual differences in baseline switching frequency.

To further decompose this index, we quantified switching and repeating behavior separately for rewarded and unrewarded trials. The probability of repeating a choice following reward was computed as:

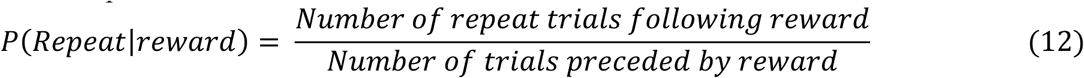

where repeat trials were defined as trials in which the same option was selected on consecutive trials. Conversely, switching following unrewarded outcomes was quantified as:

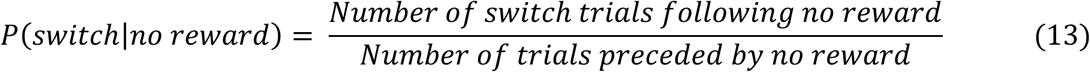

All behavioral metrics were computed at the level of individual participants or animals and subsequently averaged within species. Statistical comparisons of the learning index and outcome-dependent switching and repeating probabilities between species were performed using independent-samples *t*-tests. Effect sizes were quantified using Hedges’ *g*, and statistical significance was assessed at an alpha level of 0.05.

To assess performance across days, we calculated session-wise reward rates (rewarded trials divided by total trials) and the number of rule reversals, and tracked these measures across days.

## Author contributions

Conceptualization: Z.R. and R.C.B.

Lever Mice PRL Data acquisition: Z.R. and P.V.

Human PRL Data acquisition: R.B., M.H., and L.M.

Formal analysis: Z.R. and H.C.P.

Methodology: Z.R., E.S.I., H.C.P., and R.C.B.

Visualization: Z.R., H.C.P., and R.C.B.

Validation: Z.R. and R.C.B.

Writing—original draft: Z.R., H.C.P., and R.C.B.

Writing—review and editing: R.B.E., L.M., and R.C.B.

Funding and Resources: R.C.B.

Supervision: R.C.B.

## Data, code, and materials availability

All data and original code for data analysis will be publicly available as of the date of publication.

