## Supplementary Materials for "Shared Latent Decision Strategies Underlie Reward-Guided Behavior Across Species"

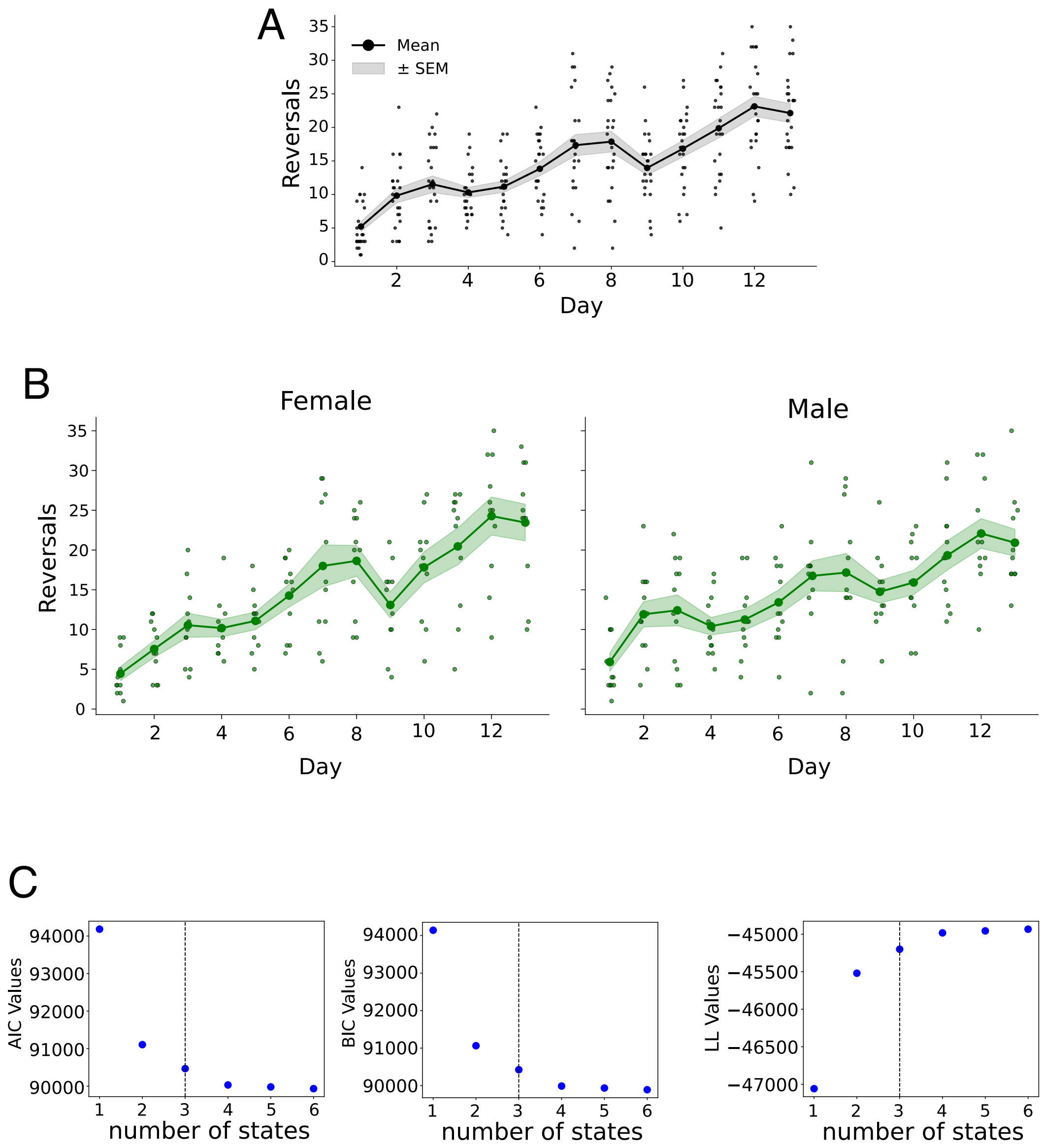


**Fig. S1. Behavioral performance and model selection in mouse probabilistic reversal learning.**
(A) Number of reversals per day across training sessions in mice. Points represent individual subjects; solid line indicates group mean; shaded region denotes SEM.
(B) Number of reversals per day separated by sex. Female (left) and male (right) mice are shown separately. Points represent individual subjects; solid line indicates group mean; shaded region denotes SEM.
(C) Model comparison metrics for GLM-HMMs with varying numbers of latent states. Akaike information criterion (AIC), Bayesian information criterion (BIC), and log-likelihood (LL) values are plotted as a function of state number. Dashed vertical line indicates the selected model.

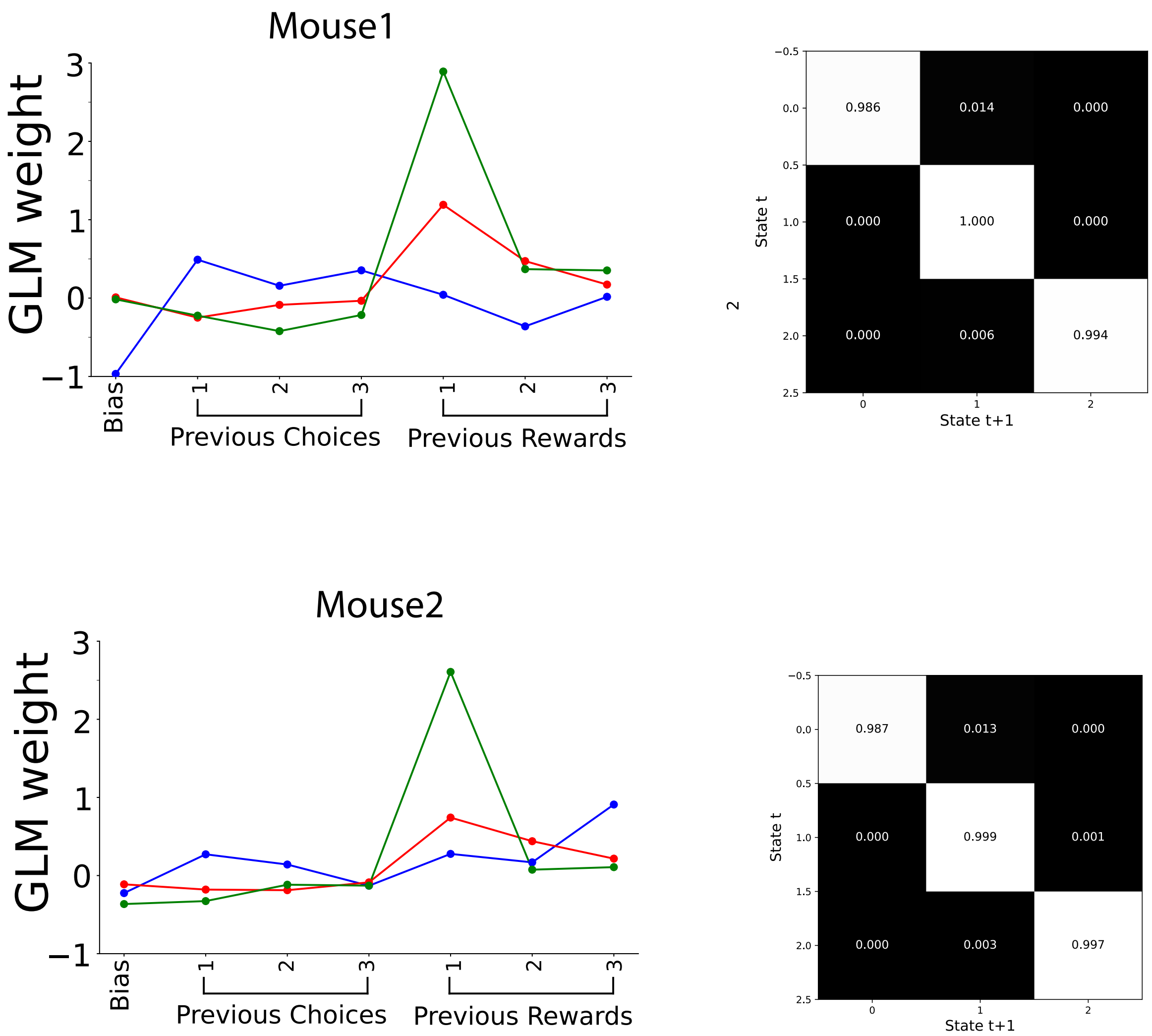


**Fig. S2. Subject-level GLM-HMM fits in mice.**
Examples of subject-level GLM-HMM fits for two representative mice. Left panels show state-specific GLM weights. Right panels show corresponding state transition matrices.

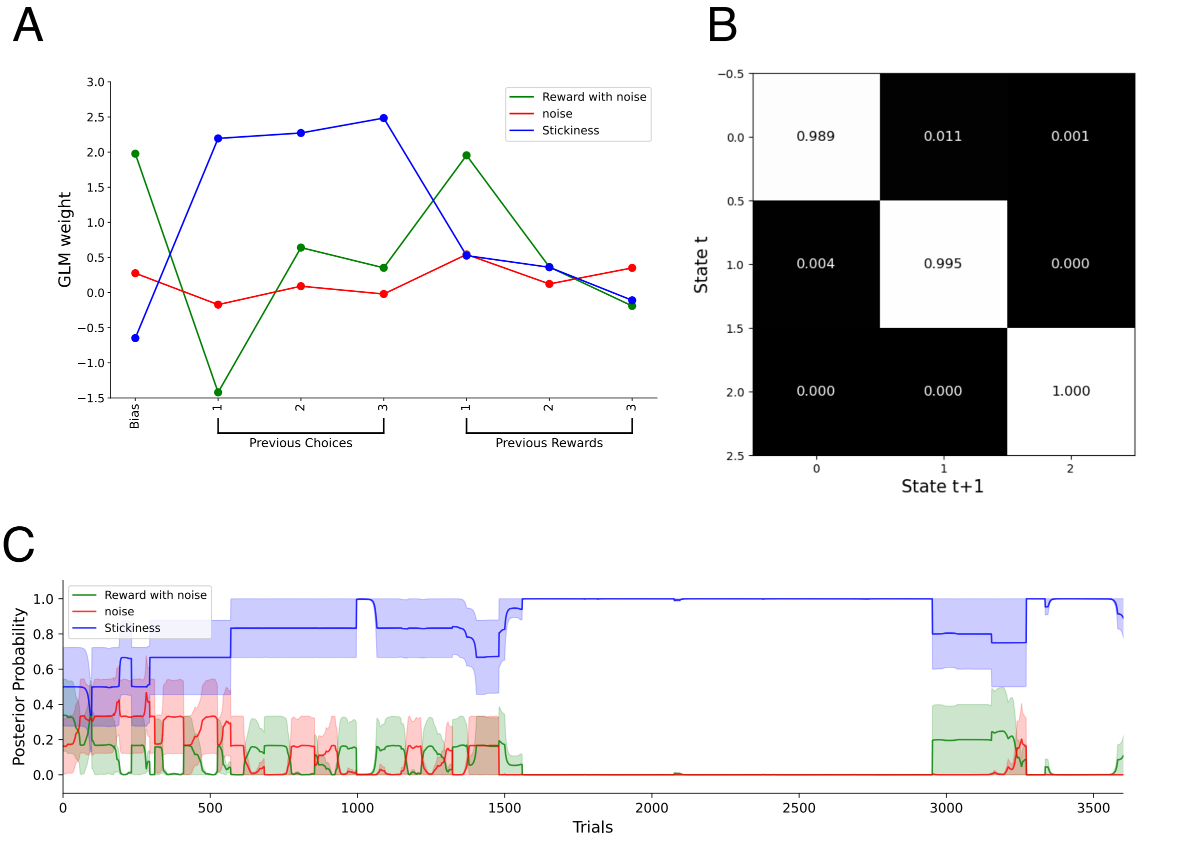


**Fig. S3. GLM-HMM fit to behavior of animals with unilateral choice.**
**(A)** State-specific GLM weights for bias, previous choices, and previous rewarded choices inferred from the GLM-HMM fit to behavior restricted to left- or right-only choices. Lines represent weights for each latent state.
**(B)** State transition matrix corresponding to the GLM-HMM fit.
**(C)** Posterior probability of latent states across trials; shaded regions denote variability across trials.

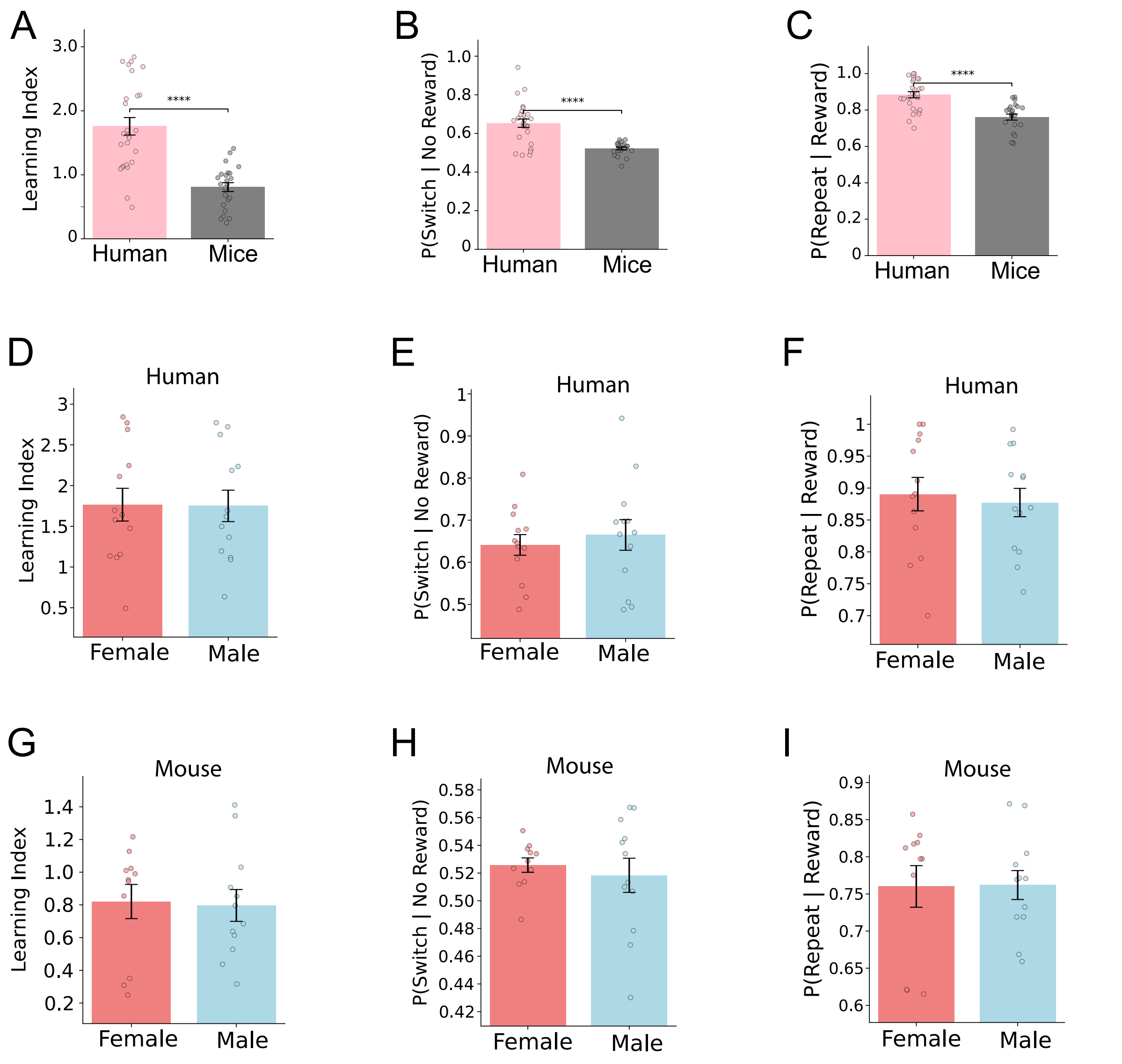


**Fig. S4. Cross-species comparison of reinforcement-dependent behavioral metrics and sex effects.**
(A–C) Comparison between humans and mice for (A) learning index, (B) probability of switching following no reward [P(switch | no reward)], and (C) probability of repeating following reward [P(repeat | reward)]. Points represent individual subjects; bars indicate mean ± SEM.
(D–F) Same measures as in (A–C) shown separately for human participants by sex: (D) learning index, (E) P(switch | no reward), and (F) P(repeat | reward). Points represent individual subjects; bars indicate mean ± SEM.
(G–I) Same measures as in (A–C) shown for mice separated by sex: (G) learning index, (H) P(switch | no reward), and (I) P(repeat | reward). Points represent individual subjects; bars indicate mean ± SEM. | ****$p < 0.0001$.

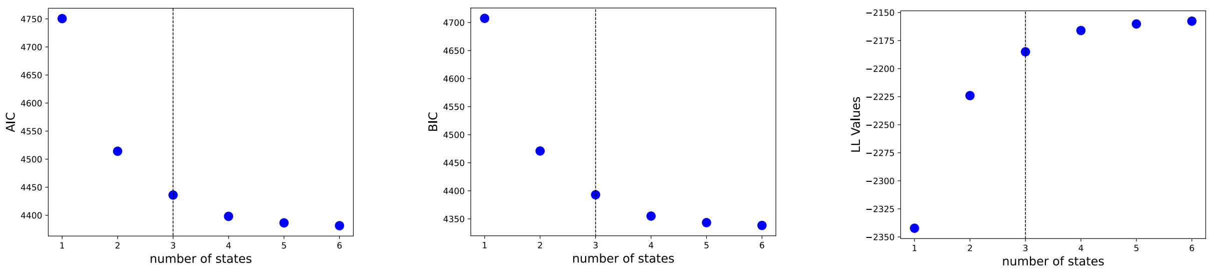


**Fig. S5. Model comparison for GLM-HMMs with varying numbers of latent states in humans.**
Akaike information criterion (AIC), Bayesian information criterion (BIC), and log-likelihood (LL) values plotted as a function of the number of latent states. Each point represents the model fit for a given state number. Dashed vertical line indicates the selected model.

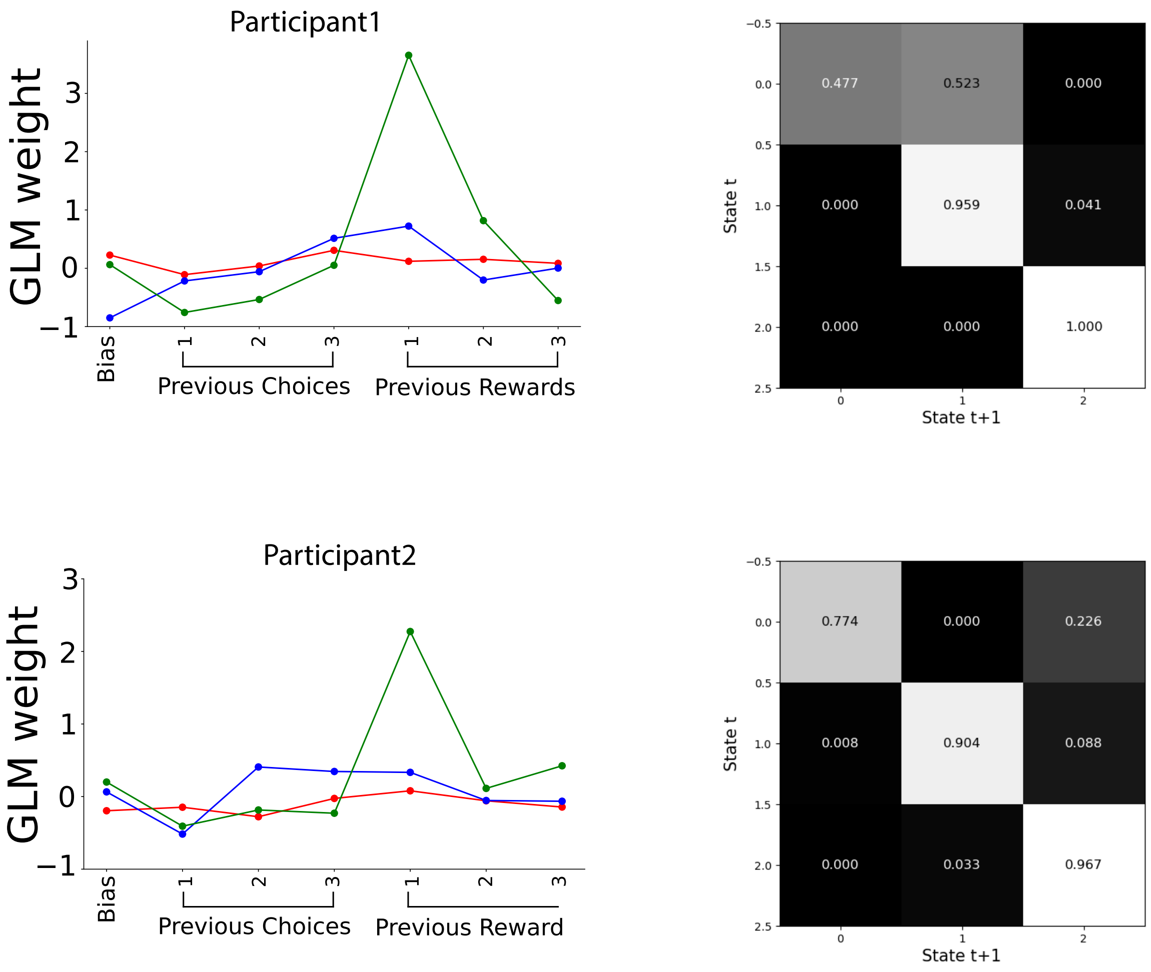


**Fig. S6. Subject-level GLM-HMM fits in human participants.**

Examples of GLM-HMM fits for two representative participants. Left panels show state-specific GLM weights for bias, previous choices, and previous rewarded choices. Right panels show the corresponding state transition matrices indicating transition probabilities between latent states.

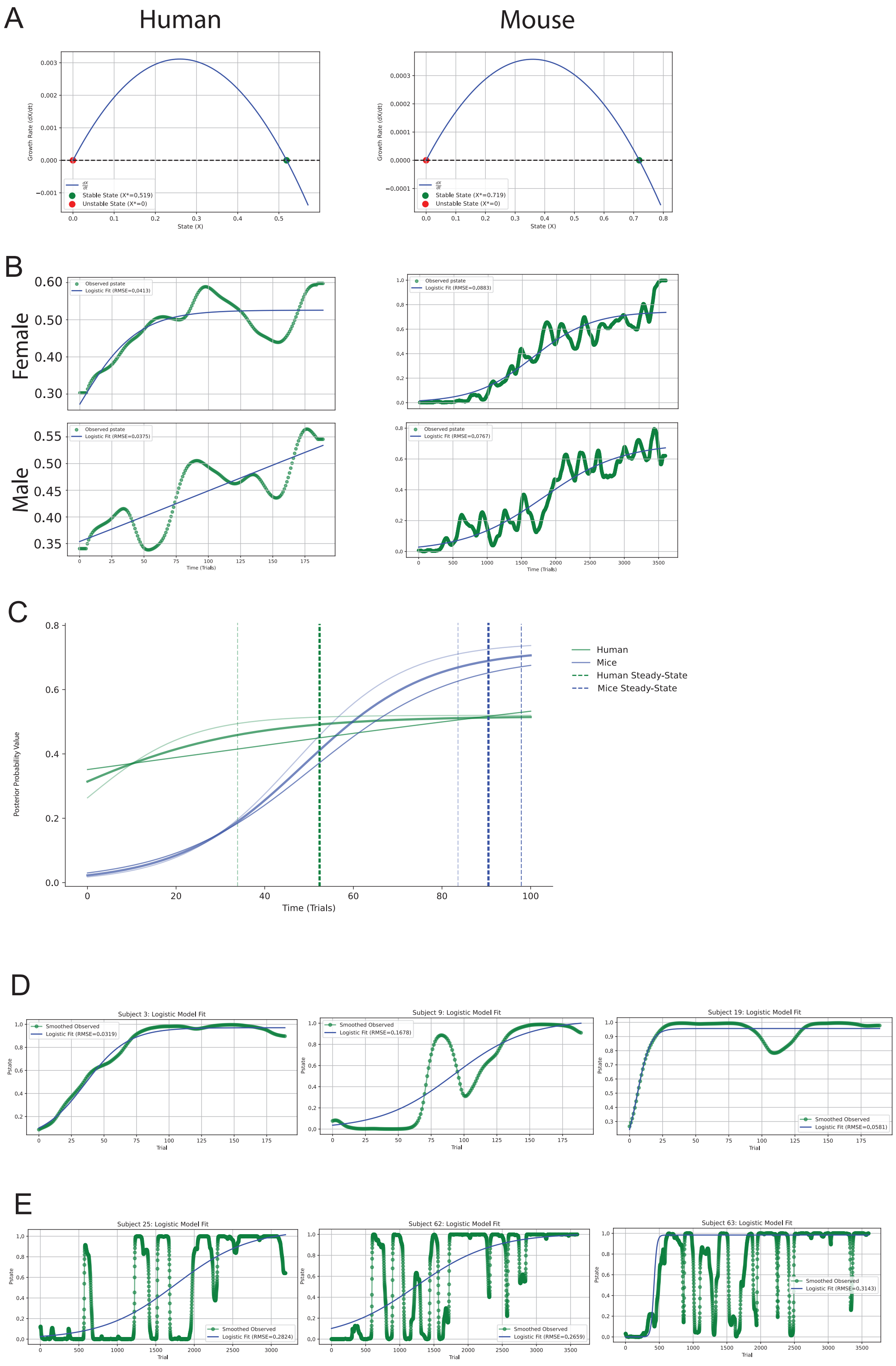
A
Human
Mouse
Growth Rate (dX/dt)
State (X)
Stable State (X*=0.519)
Unstable State (X*=0)
Stable State (X*=0.719)
Unstable State (X*=0)
B
Female
Male
Observed pstate
Logistic Fit (RMSE=0.0413)
Logistic Fit (RMSE=0.0883)
Logistic Fit (RMSE=0.0375)
Logistic Fit (RMSE=0.0767)
Time (Trials)
C
Human
Mice
Human Steady-State
Mice Steady-State
Posterior Probability Value
Time (Trials)
D
Subject 3: Logistic Model Fit
Subject 9: Logistic Model Fit
Subject 19: Logistic Model Fit
Smoothed Observed
Logistic Fit (RMSE=0.0319)
Logistic Fit (RMSE=0.1678)
Logistic Fit (RMSE=0.0581)
Pstate
Trial
E
Subject 25: Logistic Model Fit
Subject 62: Logistic Model Fit
Subject 63: Logistic Model Fit
Smoothed Observed
Logistic Fit (RMSE=0.2824)
Logistic Fit (RMSE=0.2659)
Logistic Fit (RMSE=0.3143)
Pstate
Trial

**Fig. S7. Logistic growth dynamics of the reward-learning state across species.**

(A) Phase portraits of the logistic growth model for reward-learning state probability in humans (left) and mice (right), showing stable and unstable fixed points.

(B) Logistic growth model fits to posterior probability of the reward-learning state across trials, separated by sex in humans (left) and mice (right). Observed trajectories and fitted curves are shown for female (top) and male (bottom) subjects.

(C) Joint logistic growth model fits for humans and mice. Curves represent fitted trajectories; dashed vertical lines indicate estimated steady-state points for each species.

(D) Individual logistic model fits for three representative human participants, showing posterior probability trajectories and corresponding fitted curves.

(E) Individual logistic model fits for three representative mice, showing posterior probability trajectories and corresponding fitted curves.

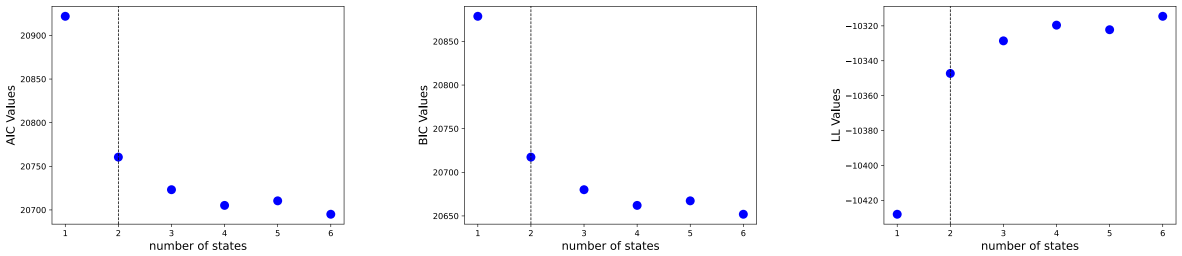


**Fig. S8. Model comparison for GLM-HMMs with varying numbers of latent states in mice touch screen PRL.**

Akaike information criterion (AIC), Bayesian information criterion (BIC), and log-likelihood (LL) values plotted as a function of the number of latent states. Each point represents the model fit for a given state number. Dashed vertical line indicates the selected model.
